# A Sparse Additive Model for High-Dimensional Interactions with an Exposure Variable

**DOI:** 10.1101/445304

**Authors:** Sahir R Bhatnagar, Tianyuan Lu, Amanda Lovato, David L Olds, Michael S Kobor, Michael J Meaney, Kieran O’Donnell, Yi Yang, Celia MT Greenwood

**Affiliations:** Department of Epidemiology, Biostatistics and Occupational Health, McGill University, Montréal, Canada; Department of Diagnostic Radiology, McGill University, Montréal, Canada; Quantitative Life Sciences, McGill University; Lady Davis Institute, Jewish General Hospital, Montréal, QC; Statistics Canada, Ottawa, ON; Department of Pediatrics, University of Colorado School of Medicine, Denver; Department of Medical Genetics, University of British Columbia, BC; Singapore Institute for Clinical Sciences, Singapore; McGill University; Department of Psychiatry, McGill University; Department of Mathematics and Statistics, McGill University; Departments of Oncology and Human Genetics, McGill University

**Keywords:** Gene-environment interaction, Strong heredity property, Blockwise coordinate descent, High-dimensional data, Variable selection

## Abstract

A conceptual paradigm for onset of a new disease is often considered to be the result of changes in entire biological networks whose states are affected by a complex interaction of genetic and environmental factors. However, when modelling a relevant phenotype as a function of high dimensional measurements, power to estimate interactions is low, the number of possible interactions could be enormous and their effects may be non-linear. In this work, we introduce a method called `sail` for detecting non-linear interactions with a key environmental or exposure variable in high-dimensional settings which respects the strong or weak heredity constraints. We prove that asymptotically, our method possesses the oracle property, i.e., it performs as well as if the true model were known in advance. We develop a computationally efficient fitting algorithm with automatic tuning parameter selection, which scales to high-dimensional datasets. Through an extensive simulation study, we show that `sail` outperforms existing penalized regression methods in terms of prediction accuracy and support recovery when there are non-linear interactions with an exposure variable. We apply `sail` to detect non-linear interactions between genes and a prenatal psychosocial intervention program on cognitive performance in children at 4 years of age. Results show that individuals who are genetically predisposed to lower educational attainment are those who stand to benefit the most from the intervention. Our algorithms are implemented in an R package available on CRAN (https://cran.r-project.org/package=sail).

## 1. Introduction

Computational approaches to variable selection have become increasingly important with the advent of high-throughput technologies in genomics and brain imaging studies, where the data has become massive, yet where it is believed that the number of truly important variables is small relative to the total number of variables. Although many approaches have been developed for main effects, there is an enduring interest in powerful methods for estimating interactions, since interactions may reflect important modulation of a genomic system by an external factor and vice versa (Bhatnagar, Yang, Khundrakpam, Evans, Blanchette, Bouchard and Greenwood, 2018).

Interactions may occur in numerous types and of varying complexities. In this paper, we consider one specific type of interaction model, where one exposure variable *E* is involved in possibly non-linear interactions with a high-dimensional set of measures **X** leading to effects on a response variable, *Y*. We propose a multivariable penalization procedure for detecting non-linear interactions between **X** and *E*. Our method is motivated by the Nurse Family Partnership (NFP); a program of prenatal and infancy home visiting by nurses for low-income mothers and their children (Olds, Henderson Jr, Cole, Eckenrode, Kitzman, Luckey, Pettitt, Sidora, Morris and Powers, 1998). In this intervention, NFP nurses guided pregnant women and parents of young children to improve the outcomes of pregnancy, their children’s health and development, and their economic self-sufficiency, with the goal of reducing disparities over the life-course. Early intervention in young children has been shown to positively impact intellectual abilities (Campbell and Ramey, 1994), and more recent studies have shown that cognitive performance is also strongly influenced by genetic factors (Rietveld, Medland, Derringer, Yang, Esko, Martin, Westra, Shakhbazov, Abdellaoui, Agrawal et al., 2013). Given the important role of both environment and genetics, we are interested in finding interactions between these two components on cognitive function in children.

### 1.1. A sparse additive interaction model

Let 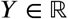 be a continuous outcome variable, 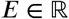 a binary or continous environment/exposure vector of known importance, and 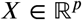 a vector of additional predictors, possibly high-dimensional. Assume that we have *n* observations of each quantity denoted by 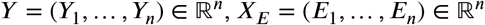, and 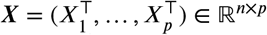. Furthermore let *f_j_*: 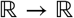 be a smoothing method for variable *X_j_* by a projection on to a set of basis functions:

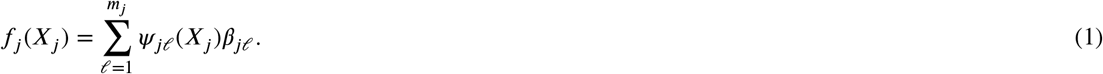

Here, the 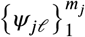 are a family of basis functions in *X_j_* (Hastie, Tibshirani and Wainwright, 2015). Let **Ψ***_j_* be the *n*×*m_j_* matrix of evaluations of the *ψ_jℓ_* and 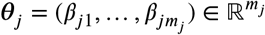 for *j* = 1,…, *p* (***θ**_j_* is a *m_j_*-dimensional column vector of basis coefficients for the *j* th main effect). In this article we consider an additive interaction regression model of the form

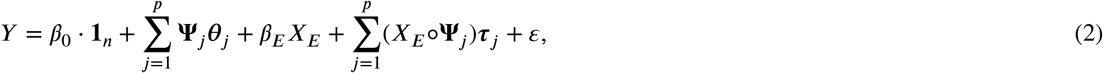

where 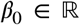 is the intercept, 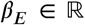 is the coefficient for the environment variable, 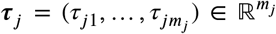 are the basis coefficients for the *j*th interaction term, (*X_E_*◦**ψ***j*) is the *n* × *m_j_* matrix formed by the component-wise multiplication of the column vector *X_E_* by each column of **Ψ***_j_*, and 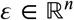 is a vector of i.i.d errors with mean zero and finite variance. Here we assume that *p* is large relative to *n*, and particularly that 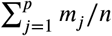 is large. Due to the large number of parameters to estimate with respect to the number of observations, one commonly-used approach in the penalization literature is to shrink the regression coefficients by placing a constraint on the values of (*β_E_, **θ**_j_, **τ**_j_*). Certain constraints have the added benefit of producing a sparse model in the sense that many of the coefficients will be set exactly to 0 (Bühlmann and Van De Geer, 2011). Such a reduced predictor set can lead to a more interpretable model with smaller prediction variance, albeit at the cost of having biased parameter estimates (Fan, Han and Liu, 2014). In light of these goals, consider the following penalized objective function:

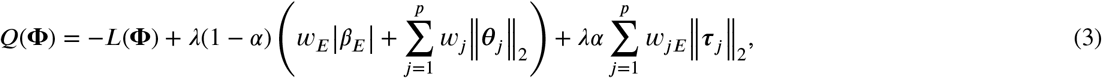

where **Φ** = (*β*_0_,*β_E_*, **θ**_1_,…, ***θ**_p_, **τ***_1_,…, ***τ**_p_*), *L*(**Φ**) is the log-likelihood function of the observations ***V**_i_* = (*Y_i_*,**Ψ***_i_*,*X_iE_*) for 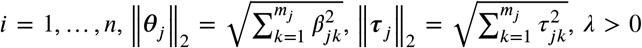 and *α* ∈ (0, 1) are adjustable tuning parameters, *w_E_, w_j_, w_jE_* are non-negative penalty factors for *j* = 1…, *p* which serve as a way of allowing parameters to be penalized differently (see Algorithm 2 for more details on how to estimate these weights). The first term in the penalty penalizes the main effects while the second term penalizes the interactions. The parameter *α* controls the relative weight on the two penalties. Note that we do not penalize the intercept.

An issue with (3) is that since no constraint is placed on the structure of the model, it is possible that an estimated interaction term is non-zero while the corresponding main effects are zero. While there may be certain situations where this is plausible, statisticians have generally argued that interactions should only be included if the corresponding main effects are also in the model (McCullagh and Nelder, 1989). This is known as the strong heredity principle (Chipman, 1996). Indeed, large main effects are more likely to lead to detectable interactions (Cox, 1984). In the next section we discuss how a simple reparametrization of the model (3) can lead to this desirable property.

### 1.2. Strong and weak heredity

The strong heredity principle states that an interaction term can only have a non-zero estimate if its corresponding main effects are estimated to be non-zero, whereas the weak heredity principle allows for a non-zero interaction estimate as long as one of the corresponding main effects is estimated to be non-zero (Chipman, 1996). In the context of penalized regression methods, these principles can be formulated as structured sparsity (Bach, Jenatton, Mairal, Obozinski et al., 2012) problems. Several authors have proposed to modify the type of penalty in order to achieve the heredity principle (Radchenko and James, 2010; Bien, Taylor, Tibshirani et al., 2013; Lim and Hastie, 2015; Haris, Witten and Simon, 2016). We take an alternative approach. Following Choi et al. (Choi, Li and Zhu, 2010), we introduce a new set of parameters 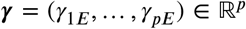 and reparametrize the coefficients for the interaction terms ***τ**_j_* in (2) as a function of *γ_jE_* and the main effect parameters ***θ**_j_* and *β_E_*. This reparametrization for both strong and weak heredity is summarized in Table 1.

**Table 1.**
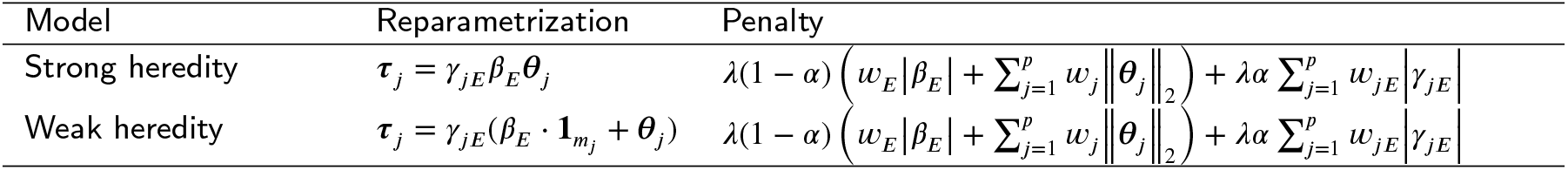
Summary of reparametrization and penalty terms for strong and weak heredity `sail` model. Note that the penalty terms are identical for both model types, i.e., the reparametrization only affects the likelihood term of the objective function.

To perform variable selection in this new parametrization, we penalize ***γ*** = (*γ*_1*E*_,…, *γ_pE_*) instead of penalizing ***τ*** as in (3), leading to the following penalized objective function:

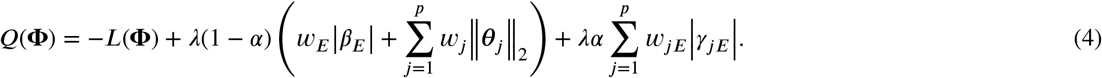

An estimate of the regression parameters is given by 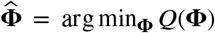. This penalty allows for the possibility of excluding the interaction term from the model even if the corresponding main effects are non-zero. Furthermore, smaller values for *α* would lead to more interactions being included in the final model while values approaching 1 would favor main effects. Similar to the elastic net (Zou and Zhang, 2009), we fix *α* and obtain a solution path over a sequence of *λ* values.

### 1.3. Toy example

We present here a toy example to better illustrate the methods proposed in this paper. With a sample size of *n* = 100, we sample *p* = 20 covariates *X*_1_,… *X_p_* independently from a *N*(0,1) distribution truncated to the interval [0,1]. Data were generated from a model which follows the strong heredity principle, but where only one covariate, *X*_2_, is involved in an interaction with a binary exposure variable (*E*):

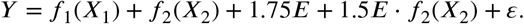

For illustration, function *f*_1_(·) is assumed to be linear, whereas function *f*_2_(·) is non-linear: *f*_1_(*x*) = −3*x*, *f*_2_(*x*) = 2(2*x* - 1)^3^. The error term *ε* is generated from a normal distribution with variance chosen such that the signal-to-noise ratio (SNR) is 2. We generated a single simulated dataset and used the strong heredity `sail` method (described below) with B-splines (df=5) to estimate the functional forms. 10-fold cross-validation (CV) was used to choose the optimal value of penalization. We used *α* = 0.5 and default values for all other arguments. We plot the solution path for both main effects and interactions in Figure 1 (top panel), coloring lines to correspond to the selected model. We see that our method is able to correctly identify the true model. We can also visually see the effect of the penalty and strong heredity principle working in tandem, i.e., the interaction term *E* · *f*_2_(*X*_2_) (orange lines in the bottom panel) can only be non-zero if the main effects *E* and *f*_2_(*X*_2_) (black and orange lines respectively in the top panel) are non-zero, while non-zero main effects does not imply a non-zero interaction.

**Figure 1:**
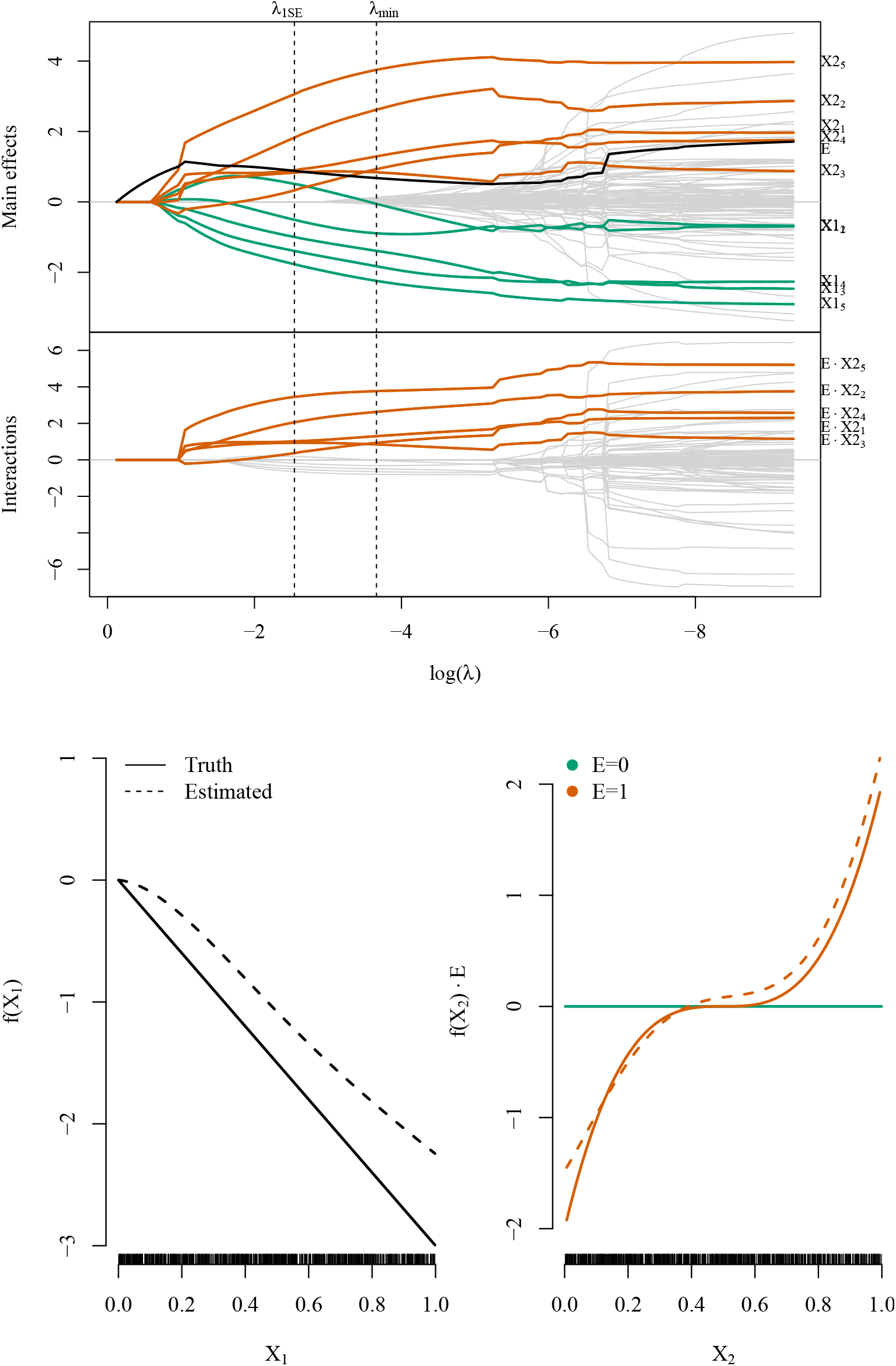
**Top**: Toy example solution path for main effects (top) and interactions (bottom). [*X*1_1_,*X*1_2_,*X*1_3_,*X*1_4_,*X*1_5_} and {*X*2_1_,*X*2_2_,*X*2_3_,*X*2_4_,*X*2_5_} are the five basis coefficients for *X*_1_ and *X*_2_, respectively. *λ*_1*SE*_ is the largest value of penalization for which the CV error is within one standard error of the minimizing value *λ_min_*. **Bottom**: Estimated smooth functions for *X*_1_ and the *X*_2_ · *E* interaction by the `sail` method based on *λ_min_*.

In Figure 1 (bottom panel), we plot the true and estimated component functions 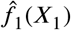 and 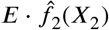, and their estimates from this analysis with `sail`. We are able to capture the shape of the correct functional form. Lack-of-fit for *f*_1_(*X*_1_) can be partially explained by acknowledging that `sail` is trying to fit a spline to a linear function. Nevertheless, this example demonstrates that `sail` can still identify trends reasonably well.

### 1.4. Related work

Methods for variable selection of interactions can be broken down into two categories: linear and non-linear interaction effects. Many of the linear effect methods consider all pairwise interactions in ***X*** (Zhao, Rocha and Yu, 2009; Choi et al., 2010; Bien et al., 2013; She and Jiang, 2014) which can be computationally prohibitive when *p* is large. More recent proposals for selection of interactions allow the user to restrict the search space to interaction candidates (Lim and Hastie, 2015; Haris et al., 2016). This is useful when the researcher wants to impose prior information on the model. Two-stage procedures, where interaction candidates are considered from an original screen of main effects, have shown good performance when *p* is large (Hao, Feng and Zhang, 2018; Shah, 2016) in the linear setting. There are many fewer methods available for estimating non-linear interactions. For example, Radchenko and James (2010) (Radchenko and James, 2010) proposed a model of the form 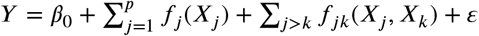, where *f*(·) are smooth component functions. This method is more computationally expensive than `sail` since it considers all pairwise interactions between the basis functions, and its effectiveness in simulations or real-data applications is unknown as there is no software implementation.

The main contributions of this paper are five-fold. First, we develop a model for non-linear interactions with a key exposure variable, following either the weak or strong heredity principle, that is computationally efficient and scales to the high-dimensional setting (*n* ≪ *p*). Second, through simulation studies, we show improved performance in terms of prediction accuracy and support recovery over existing methods that only consider linear interactions or additive main effects. Third, we show that our method possesses the oracle property (Fan and Li, 2001), i.e., it performs as well as if the true model were known in advance. Fourth, we demonstrate the performance of our method in two applications: 1) gene-environment interactions in a prenatal psychosocial intervention program Olds et al. (1998) and 2) a study aimed at identifying which clinical variables influence mortality rates amongst seriously ill hospitalized patients (Connors, Dawson, Desbiens, Fulkerson, Goldman, Knaus, Lynn, Oye, Bergner, Damiano et al., 1995). Fifth, we implement our algorithms in the `sail` R package on CRAN (https://cran.r-project.org/package=sail), along with extensive documentation. In particular, our implementation also allows for linear interaction models, user-defined basis expansions, a cross-validation procedure for selecting the optimal tuning parameter, and differential shrinkage parameters to apply the adaptive lasso idea (Zou, 2006).

The rest of the paper is organized as follows. Section 2 describes our optimization procedure and some details about the algorithm used to fit the `sail` model for the least squares case. Theoretical results are given in Section 3. In Section 4, through simulation studies we compare the performance of our proposed approach and demonstrate the scenarios where it can be advantageous to use `sail` over existing methods. Section 5 contains two real data examples and Section 6 discusses some limitations and future directions.

## 2. Computation

In this section we describe a blockwise coordinate descent algorithm for fitting the least-squares version of the `sail` model in (4). We fix the value for *α* and minimize the objective function over a decreasing sequence of *λ* values (*λ_max_* >···> *λ_min_*). We use the subgradient equations to determine the maximal value *λ_max_* such that all estimates are zero. Due to the heredity principle, this reduces to finding the largest *λ* such that all main effects (*β_E_, **θ***_1_,…, ***θ**_p_*) are zero. Following Friedman et al. (Friedman, Hastie and Tibshirani, 2010), we construct a *λ*-sequence of 100 values decreasing from *λ_max_* to 0.001 *λ_max_* on the log scale, and use the warm start strategy where the solution for *λ_ℓ_* is used as a starting value for *λ*_*ℓ*+1_.

### 2.1. Blockwise coordinate descent for least-squares loss

We assume that *Y*, **Ψ***_j_*, *X_E_* and *X_E_*◦**ψ***_j_* have been centered by their sample means 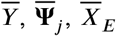, and 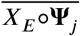, respectively. Here, 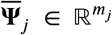 and 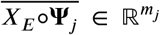 represent the column means of **Ψ***_j_* and *X_E_* ◦**Ψ***_j_*, respectively. Since the intercept (*β*_0_) is not penalized and all variables have been centered, we can omit it from the loss function and compute it once the algorithm has converged for all other parameters. The strong heredity `sail` model with least-squares loss has the form:

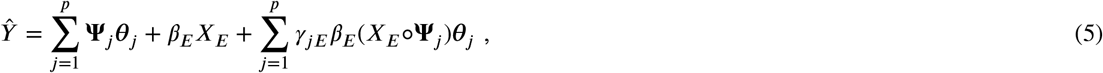

and the objective function is given by

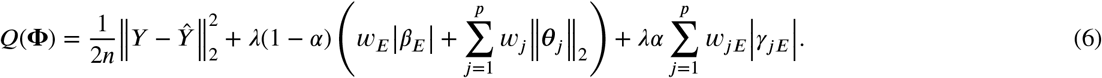

Solving (6) in a blockwise manner allows us to leverage computationally fast algorithms for *ℓ*_1_ and *ℓ*_2_ norm penalized regression. Indeed, by careful construction of pseudo responses and pseudo design matrices, existing efficient algorithms can be used to estimate the parameters. The objective function simplifies to a modified lasso problem when holding all *θ_j_* fixed, and a modified group lasso problem when holding *β_E_* and all *γ_jE_* fixed. The main computations are provided in Algorithm 1. A more detailed version of the derivations are given in Supplemental Section B.1.

#### Algorithm 1 Blockwise Coordinate Descent for Least-Squares sail with Strong Heredity

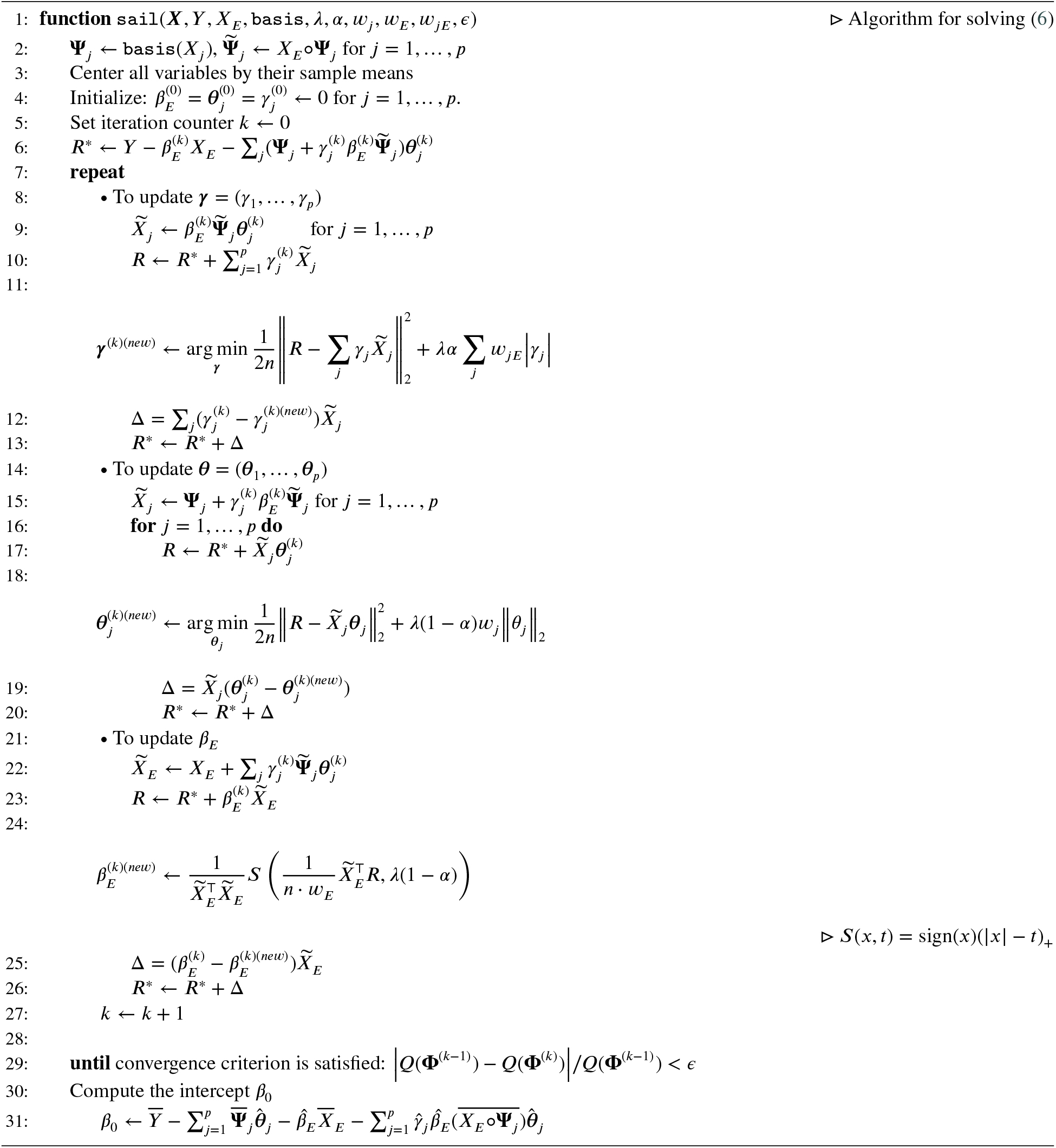

### 2.2 Details on Update for *θ*

Here we discuss a computational speedup in the updates for the ***θ*** parameter. The partial residual (*R_s_*) used for updating ***θ**_s_* (*s* ∈ 1,…,*p*) at the *k*th iteration is given by

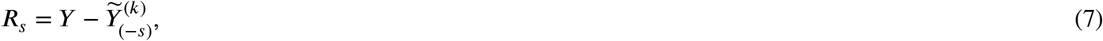

where 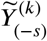 is the fitted value at the *k*th iteration excluding the contribution from **Ψ**_*s*_:

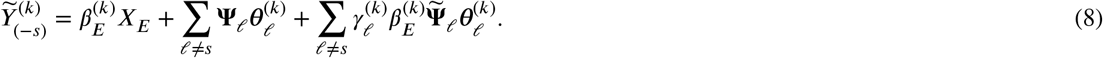

Using (8), (7) can be re-written as

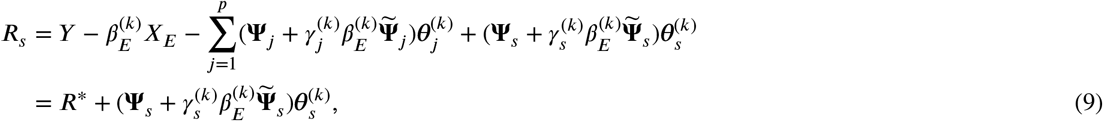

where

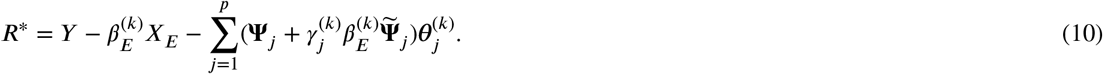

Denote 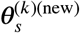 the solution for predictor *s* at the *k*th iteration, given by:

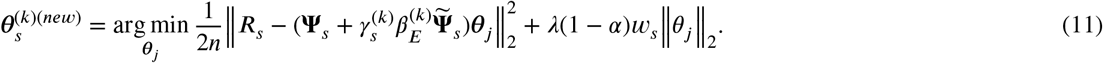

Now we want to update the parameters for the next predictor ***θ***_*s*+1_ (*s* + 1 ∈ 1,…,*p*) at the *k*th iteration. The partial residual used to update ***θ***_*s*+1_ is given by

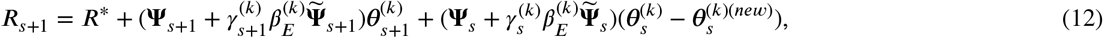

where *R*\* is given by (10), 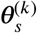 is the parameter value prior to the update, and 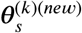 is the updated value given by (11). Taking the difference between (9) and (12) gives

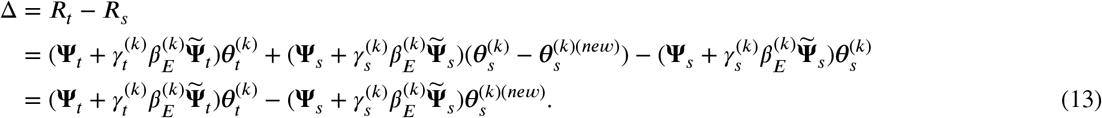

Therefore *R_t_* = *R_s_* + Δ, and the partial residual for updating the next predictor can be computed by updating the previous partial residual by Δ, given by (13). This formulation can lead to computational speedups especially when Δ = 0, meaning the partial residual does not need to be re-calculated.

### 2.3. Weak Heredity

Our method can be easily adapted to enforce the weak heredity property. That is, an interaction term can only be present if at least one of its corresponding main effects is non-zero. To do so, we reparametrize the coefficients for the interaction terms in (2) as ***τ**_j_* = *γ_jE_*(*β_E_* · **1***_m_j__* + ***θ**_j_*), where **1***_m_j__* is a vector of ones with dimension *m_j_* (i.e. the length of ***θ**_j_*). We defer the algorithm details for fitting the `sail` model with weak heredity in Supplemental Section B.4, as it is very similar to Algorithm 1 for the strong heredity `sail` model.

### 2.4. Adaptive sail

The weights for the environment variable, main effects and interactions are given by *w_E_, w_j_* and *w_jE_* respectively. These weights serve as a means of allowing a different penalty to be applied to each variable. In particular, any variable with a weight of zero is not penalized at all. This feature is usually selected for one of two reasons:

1. Prior knowledge about the importance of certain variables is known. Larger weights will penalize the variable more, while smaller weights will penalize the variable less
2. Allows users to apply the adaptive `sail`, similar to the adaptive lasso (Zou, 2006)

We describe the adaptive `sail` in Algorithm 2. This is a general procedure that can be applied to the weak and strong heredity settings. We provide this capability in the `sail` package using the `penalty.factor` argument.

#### Algorithm 2 Adaptive sail algorithm

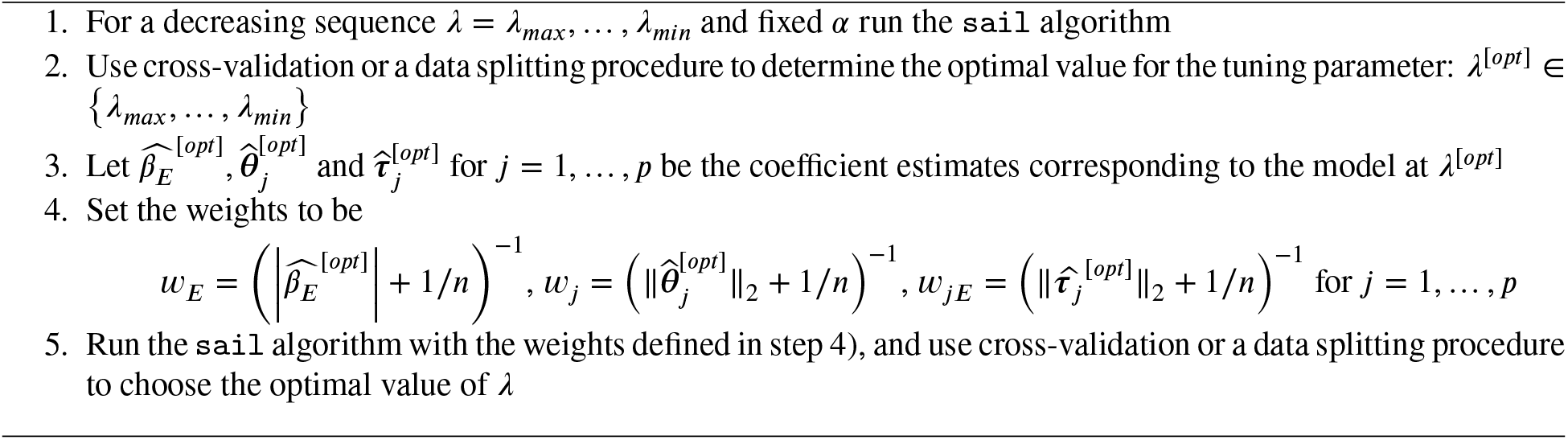

### 2.5. Flexible design matrix

The definition of the basis expansion functions in (1) is very flexible, in the sense that our algorithms are independent of this choice. As a result, the user can apply any basis expansion they desire. In the extreme case, one could apply the identity map, i.e., *f_j_*(*X_j_*) = *X_j_* which leads to a linear interaction model (referred to as `linear sail`). When little information is known a priori about the relationship between the predictors and the response, by default, we choose to apply the same basis expansion to all columns of ***X***. This is a reasonable approach when all the variables are continuous. However, there are often situations when the data contains a combination of categorical and continuous variables. In these cases it may be sub-optimal to apply a basis expansion to the categorical variables. Owing to the flexible nature of our algorithm, we can handle this scenario in our implementation by allowing a user-defined design matrix. The only extra information needed is the group membership of each column in the design matrix. We illustrate such an example in a vignette of the `sail R` package.

## 3. Theory

In this section we study the asymptotic behaviour of the `sail` estimator 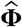, defined as the minimizer of (4), as well as the model selection properties. We show that `sail` possesses the oracle property when the sample size approaches infinity and the number of predictors is fixed. That is, under certain regularity conditions, it performs as well as if the true model were known in advance and has the optimal estimation rate (Zou, 2006). The regularity conditions and proofs are given in Supplemental Section 1.

Let 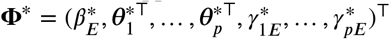 denote the unknown vector of true coefficients in (4). To simplify the notation, we use the representation 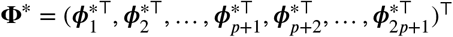, where 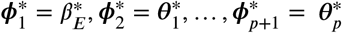, and 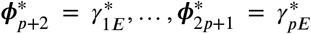. Denote by 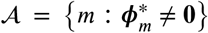 the unknown sparsity pattern of **Φ**\*, and 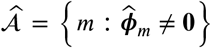 the estimated `sail` model selector. We can rewrite the penalty terms in (4), and consider the `sail` estimates 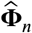 given b

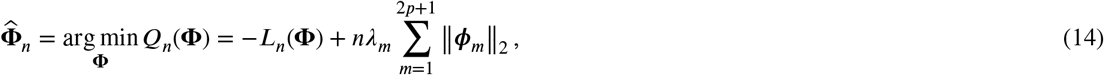

where *λ*_1_ = *λ*(1 - *α*)*w_E_, λ_m_* = *λ*(1 - *α*)*w_m_* for *m* = 2,…, *p* + 1 and *λ_m_* = *λαw_mE_* for *m* = *p* + 2,…, 2*p* + 1. Define

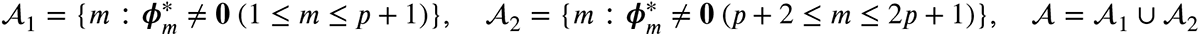

that is, 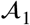 contains the indices for main effects whose true coefficients are non-zero, and 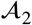 contains the indices for interaction terms whose true coefficients are non-zero. Let

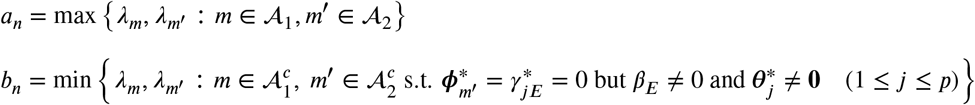

Note that our asymptotic results are stated for the main effects and interaction terms only, even though our formulation includes an unpenalized intercept. Consistency results immediately follow for *β*_0_ since we assume the data has been centered, leading to a closed form solution for the intercept in the least-squares setting.

### Lemma 1.

*[Existence of a local minimizer] If 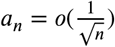 as n → ∞, i.e. 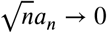 then 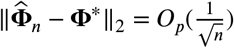*

Lemma (1) states that if the tuning parameters corresponding to the non-zero coefficients converge to 0 at a speed faster than 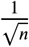, then there exists a local minimizer of *Q_n_*(**Φ**) which is 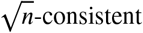 (Wang, Li and Tsai, 2007; Choi et al., 2010).

### Theorem 1

(Model selection consistency). *If 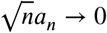 and 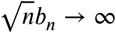, then*

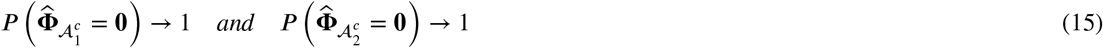

Theorem (1) shows that `sail` can consistently remove the main effects and interaction terms which are not associated with the response with high probability. Together with Lemma (1), we see that the asymptotic behaviour of the penalty terms for the zero and non-zero predictors must be different to satisfy the model selection consistency property (15) (Nardi, Rinaldo et al., 2008). Specifically, when the tuning parameters for the non-zero coefficients converge to 0 faster than 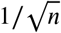 (i.e. 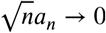) and those for zero coefficients are large enough (i.e. 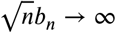), the Lemma (1) and Theorem (1) imply that the 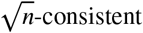 estimator 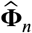 satisfies 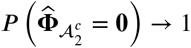.

Next, we obtain the asymptotic distribution of the `sail` estimator.

### Theorem 2

(Asymptotic normality). *Denote 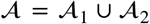. Assume that 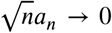 and 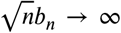. Under the regularity conditions, the subvector 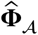 of the local minimizer 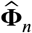 given in Lemma (1) satisfies*

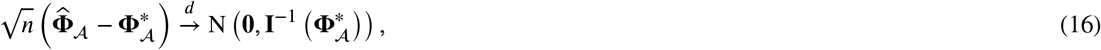

*where* 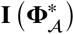 *is the Fisher information matrix for* 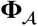 *at* 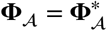, *assuming* 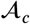 *is known in advance*.

Together, Theorems (1) and (2) establish that if the tuning parameters satisfy the conditions 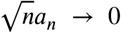 and 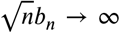, then as the sample size grows large, `sail` has the oracle property (Fan and Li, 2001). In order for the conditions on the tuning parameters to be satisfied, we follow the strategies outlined for the adaptive Lasso (Zou, 2006), the adaptive group Lasso (Nardi et al., 2008) and the adaptive elastic-net (Zou and Zhang, 2009). That is, we define the adaptive weights as 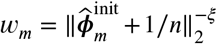 for *m* = 1,…, 2*p*+1, where *ξ* is a positive constant and 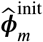 is an initial 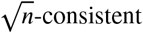 estimate of 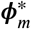. Here, the 1/*n* is to avoid division by zero.

## 4 Simulation Study

In this section, we use simulated data to understand the performance of `sail` in different scenarios.

### 4.1. Comparator Methods

Since there are no other packages that directly address our chosen problem, we selected comparator methods based on the following criteria: 1) penalized regression methods that can handle high-dimensional data (*n* < *p*), 2) allowing at least one of linear effects, non-linear effects or interaction effects, and 3) having a software implementation in R. The selected methods can be grouped into three categories:

1. Linear main effects: `lasso` (Tibshirani, 1996), `adaptive lasso` (Zou, 2006)
2. Linear interactions: `lassoBT` (Shah, 2016), `GLinternet` (Lim and Hastie, 2015)
3. Non-linear main effects: `HierBasis` (Haris, Shojaie and Simon, 2019), `SPAM` (Ravikumar, Lafferty, Liu and Wasserman, 2009), `gamsel` (Chouldechova and Hastie, 2015)

For `GLinternet` we specified the `interactioncandidates` argument so as to only consider interactions between the environment and all other *X* variables. For all other methods we supplied (***X**, X_E_*) as the data matrix, 100 for the number of tuning parameters to fit, and used the default values otherwise (`R` code for each method available at https://github.com/sahirbhatnagar/sail/blob/master/my_sims/method_functions.R). `lassoBT` considers all pairwise interactions as there is no way for the user to restrict the search space. `SPAM` applies the same basis expansion to every column of the data matrix; we chose 5 basis spline functions. `HierBasis` and `gamsel` selects whether a term in an additive model is non-zero, linear, or a non-linear spline up to a specified max degrees of freedom per variable.

We compare the above listed methods with our main proposal method `sail`, as well as with `adaptive sail` (Algorithm 2) and `sail weak` which has the weak heredity property. For each function *f_j_*, we use a B-spline basis matrix with `degree=5` implemented in the bs function in `R` (R Core Team, 2017). We center the environment variable and the basis functions before running the `sail` method.

### 4.2. Simulation Design

To make the comparisons with other methods as fair as possible, we followed a simulation framework that has been previously used for variable selection methods in additive models (Lin, Zhang et al., 2006; Huang, Horowitz and Wei, 2010). We extend this framework to include interaction effects as well. The covariates are simulated as follows. First, we generate *x*_1_,…, *x*_1000_ independently from a standard normal distribution truncated to the interval [0,1] for *i* = 1,…, *n*. The first four variables are non-zero (i.e. active in the response), while the rest of the variables are zero (i.e. are noise variables). The exposure variable (*X_E_*) is generated from a standard normal distribution truncated to the interval [-1,1]. The outcome *Y* is then generated following one of the models and assumptions described below. We evaluate the performance of our method on three of its defining characteristics: 1) the strong heredity property, 2) nonlinearity of predictor effects and 3) interactions. Simulation scenarios are designed specifically to test the performance of these characteristics.

1. **Heredity simulation**
  Scenario (a) <u>Truth obeys strong heredity</u>. In this situation, the true model for *Y* contains main effect terms for all covariates involved in interactions.

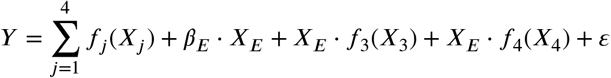
  Scenario (b) <u>Truth obeys weak heredity</u>. Here, in addition to the interaction, the *E* variable has its own main effect but the covariates *X*_3_ and *X*_4_ do not.

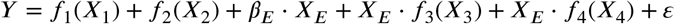
  Scenario (c) <u>Truth only has interactions</u>. In this simulation, the covariates involved in interactions do not have main effects as well.

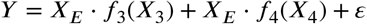
2. **Non-linearity simulation scenario**
  <u>Truth is linear</u>. `sail` is designed to model non-linearity; here we assess its performance if the true model is completely linear.

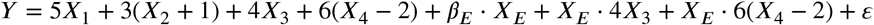
3. **Interactions simulation scenario**
  <u>Truth only has main effects</u>. `sail` is designed to capture interactions; here we assess its performance when there are none in the true model.

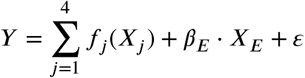

The true component functions are the same as in (Lin et al., 2006; Huang et al., 2010) and are given by *f*_1_(*t*) = 5*t*, *f*_2_(*t*) = 3(2*t* - 1)^2^, *f*_3_(*t*) = 4sin(2*πt*)/(2 - sin(2*πt*)), *f*_4_(*t*) = 6(0.1 sin(2*πt*) + 0.2cos(2*πt*) + 0.3sin(2*πt*)^2^ + 0.4cos(2*πt*)^3^ + 0.5 sin(2*πt*)^3^). We set *β_E_* =2 and draw *ε* from a normal distribution with variance chosen such that the signal-to-noise ratio is 2. Using this setup, we generated 200 replications consisting of a training set of *n* = 200, a validation set of *n* = 200 and a test set of *n* = 800. The training set was used to fit the model and the validation set was used to select the optimal tuning parameter corresponding to the minimum prediction mean squared error (MSE). Variable selection results including true positive rate, false positive rate and number of active variables (the number of variables with a non-zero coefficient estimate) were assessed on the training set, and MSE was assessed on the test set.

### 4.3. Results

The prediction accuracy and variable selection results for each of the five simulation scenarios are shown in Figure 2 and Table 2, respectively. We see that `sail, adaptive sail` and `sail weak` have the best performance in terms of both MSE and yielding correct sparse models when the truth follows a strong heredity (scenario 1a), as we would expect, since this is exactly the scenario that our method is trying to target. Our method is also competitive when only main effects are present (scenario 3) and performs just as well as methods that only consider linear and non-linear main effects (`HierBasis, SPAM`), owing to the penalization applied to the interaction parameter. Due to the heredity property being violated in scenario 1c), no method can identify the correct model with the exception of `GLinternet`. When only linear effects and interactions are present (scenario 2), we see that `adaptive sail` has similar MSE compared to the other linear interaction methods (`lassoBT` and `GLinternet`) with a better TPR and FPR. It is important to note that the variable selection performance of `sail` is highly dependent on being able to correctly select the exposure variable (*X_E_*). In Supplemental Section C, we show the selection rates of *X_E_*. We see that `sail` is able to consistently identify the exposure variable across all simulation scenarios and replications. Overall, our simulation study results suggests that `sail` outperforms existing methods when the true model contains non-linear interactions, and is competitive even when the truth only has either linear or additive main effects.

**Figure 2:**
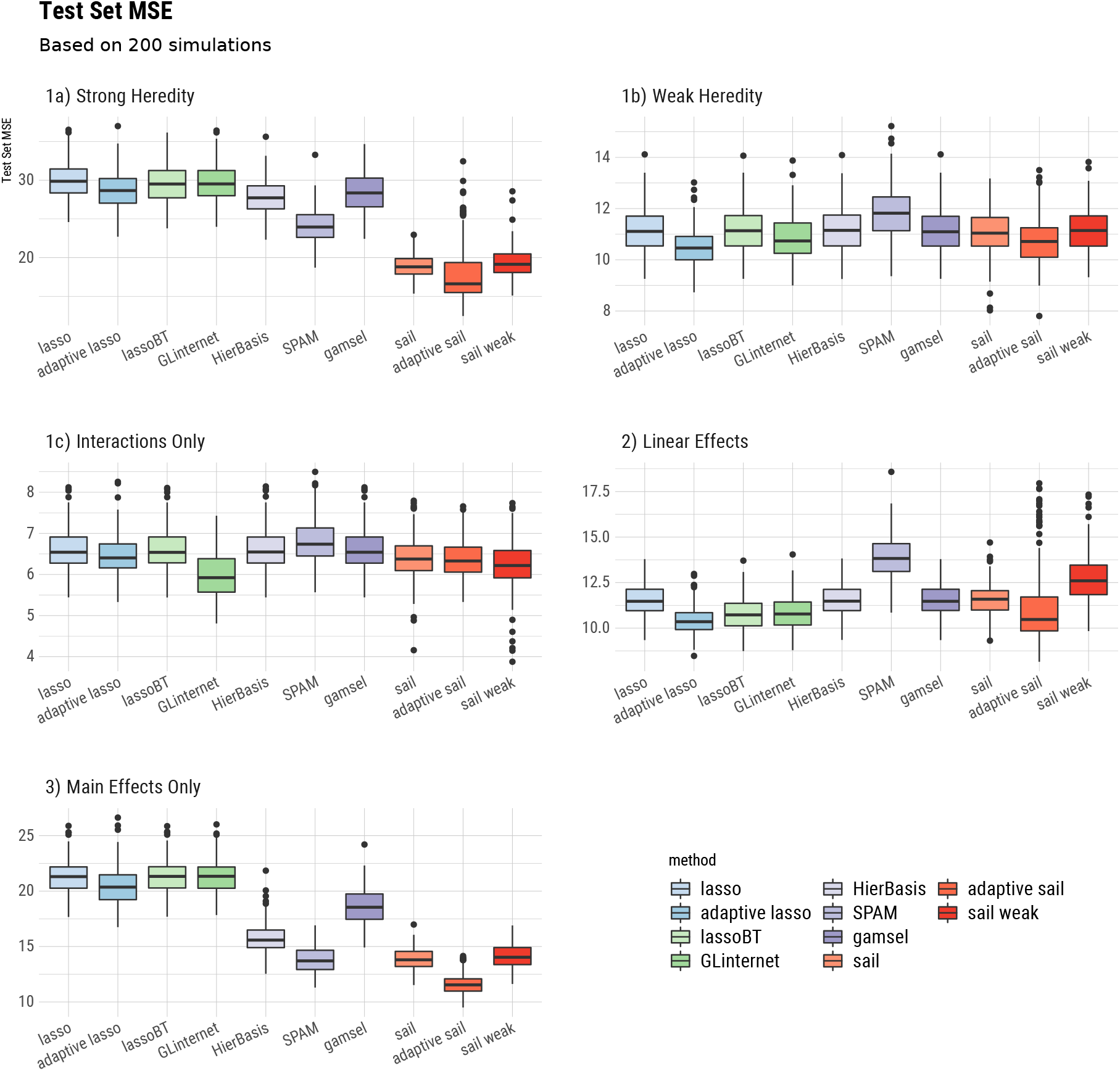
Boxplots of the test set mean squared error from 200 replications for each of the five simulation scenarios.

**Table 2:**
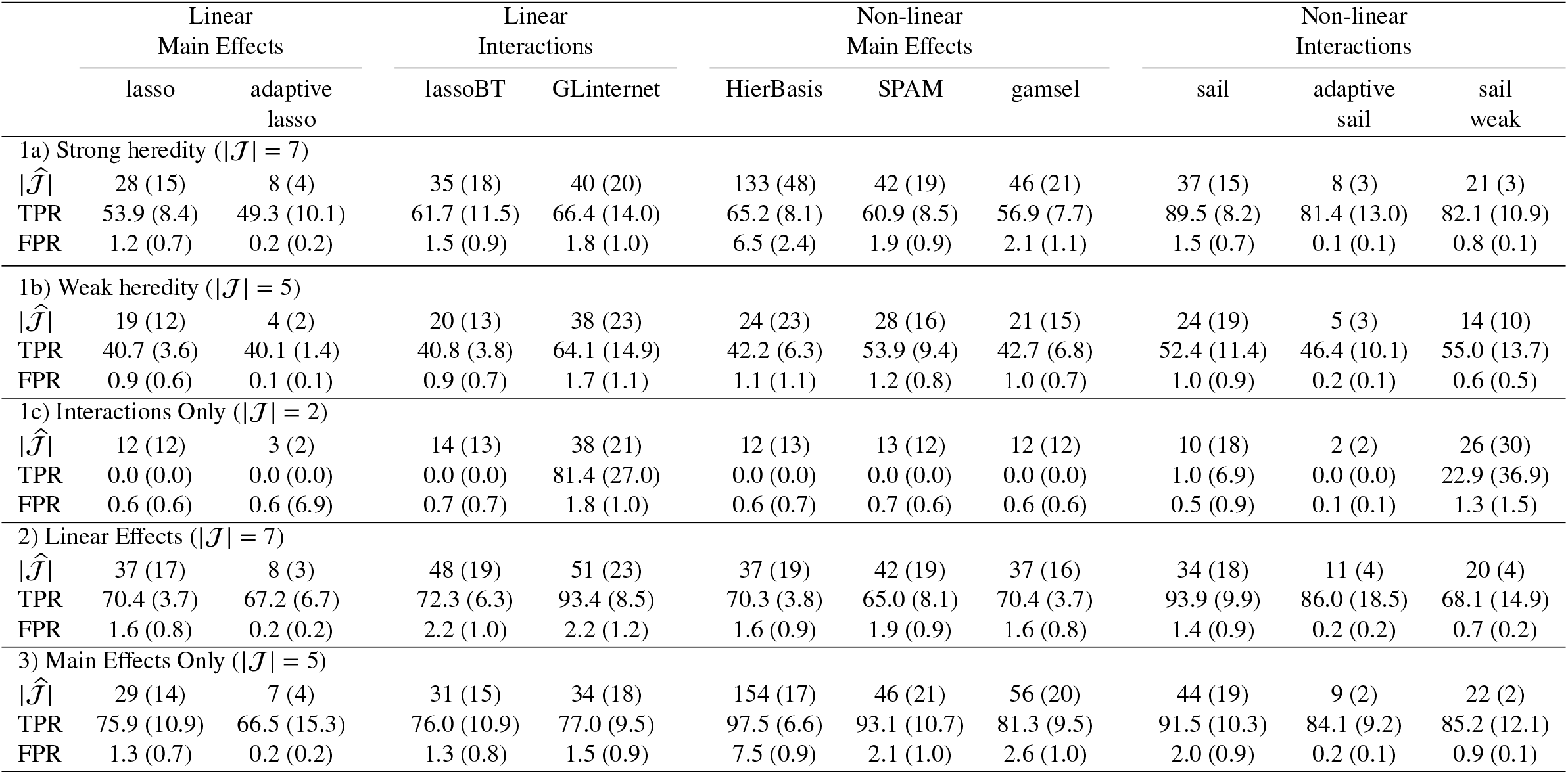
Mean (standard deviation) of the number of selected variables 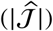, true positive rate (TPR) and false positive rate (FPR) as a percentage from 200 replications for each of the five scenarios. 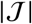 is the number of truly associated variables.

We also plotted the true and predicted curves for scenario 1a) in Supplemental Section C, to visually inspect whether our method could correctly capture the shape of the association between the predictors and the response for both main and interaction effects. In general, we see the non-linear effects are clearly being captured by `sail`.

## 5. Real data applications

### 5.1. Gene-environment interactions in the Nurse Family Partnership program

It is well known that environmental exposures can have an important impact on academic achievement. Indeed, early intervention in young children has been shown to positively impact intellectual abilities (Campbell and Ramey, 1994). More recent studies have shown that cognitive performance, a trait that measures the ability to learn, reason and solve problems, is also strongly influenced by genetic factors. Genome-wide association studies (GWAS) suggest that 20% of the variance in educational attainment (years of education) may be accounted for by common genetic variation (Rietveld et al., 2013; Okbay, Beauchamp, Fontana, Lee, Pers, Rietveld, Turley, Chen, Emilsson, Meddens et al., 2016). Unsurprisingly, there is significant overlap in the SNPs that predict educational attainment and measures of cognitive function. An interesting query that arises is how the environment interacts with these genetics variants to predict measures of cognitive function. To address this question, we analyzed data from the Nurse Family Partnership (NFP), a psychosocial intervention program that begins in pregnancy and targets maternal health, parenting and mother-infant interactions (Olds et al., 1998). The Stanford Binet IQ scores at 4 years of age were collected for 189 subjects (including 19 imputed using `mice` (Buuren and Groothuis-Oudshoorn, 2010)) born to women randomly assigned to control (*n* = 100) or nurse-visited intervention groups (*n* = 89). For each subject, we calculated a polygenic risk score (PRS) for educational attainment at different p-value thresholds using weights from the GWAS conducted in Okbay et al. (Okbay et al., 2016). In this context, individuals with a higher PRS have a propensity for higher educational attainment. The goal of this analysis was to determine if there was an interaction between genetic predisposition to educational attainment (*X*) and maternal participation in the NFP program (*E*) on child IQ at 4 years of age (*Y*). We applied the weak heredity `sail` with cubic B-splines and *α* = 0.1 to encourage interactions, and selected the optimal tuning parameter using 10-fold cross-validation. Our method identified an interaction between the intervention and PRS which included genetic variants at the 0.0001 level of significance. This interaction is shown in Figure 3. We see that the intervention has a much larger effect on IQ for lower PRS compared to a higher PRS. In other words, perinatal home visitation by nurses can impact IQ scores in children who are genetically predisposed to lower educational attainment. Similar results were obtained for the other imputed datasets (Supplemental Section D). We also compared `sail` with two other interaction selection methods, `lassoBT` and `GLinternet` with default settings, on 200 bootstrap samples of the data. The average and standard deviation of the MSE and size of the active set 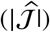 across the 200 bootstrap samples are given in Table 3. We see that `sail` tends to select sparser models while maintaining similar prediction performance compared to `lassoBT`. The `GLinternet` statistics are omitted here since the algorithm did not converge for many of the 200 simulations.

**Figure 3:**
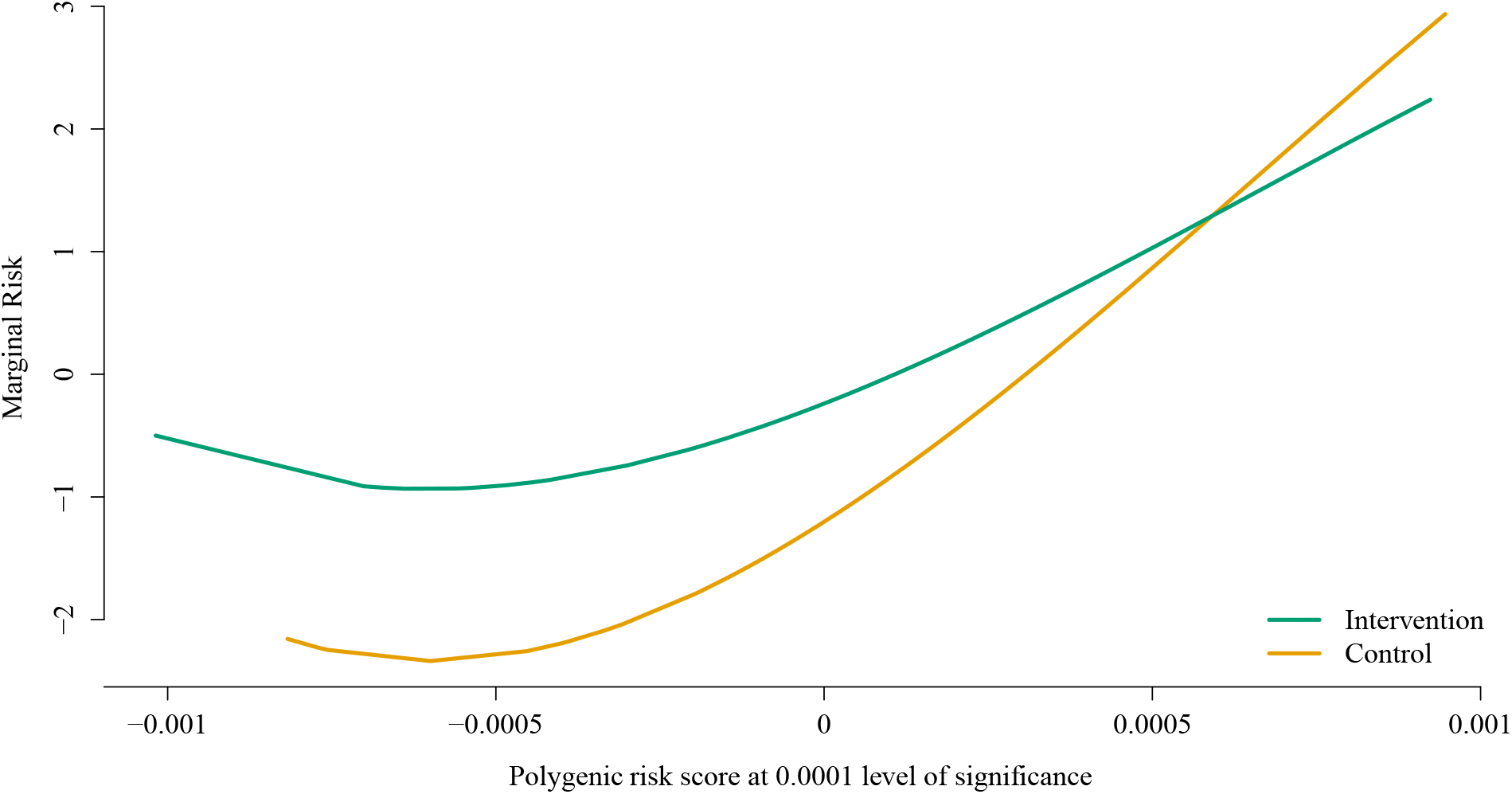
Estimated interaction effect identified by the weak heredity `sail` using cubic B-splines and *α* = 0.1 for the Nurse Family Partnership data. The selected model, chosen via 10-fold cross-validation, contained three variables: the main effects for the intervention and the PRS for educational attainment using genetic variants significant at the 0.0001 level, as well as their interaction.

**Table 3.** Comparison of analytic methods for selecting interactions using the Nurse Family Partnership program and the SUPPORT datasets. Averages (standard deviations in parentheses) are based on 200 bootstrap samples. 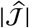 is the number of variables selected by the method. `GLinternet` results not reported for NFP data since the algorithm did not converge in many of the bootstrap samples.

| Method | Nurse Family Partnership |  | SUPPORT |  |
| --- | --- | --- | --- | --- |
| | Mean Squared Error | $\vert \hat{\mathcal{J}}\vert $ | AUC | $\vert \hat{\mathcal{H}}\vert $ |
| `sail` | 3.5 (0.6) | 4 (3) | 0.66 (0.01) | 25 (3) |
| `lassoBT` | 3.53 (0.477) | 11 (6) | 0.65 (0.009) | 49 (14) |
| `GLinternet` | — | — | 0.65 (0.009) | 58 (7) |

### 5.2. Study to Understand Prognoses Preferences Outcomes and Risks of Treatment

The Study to Understand Prognoses Preferences Outcomes and Risks of Treatment (SUPPORT) aimed at identifying which clinical variables influence medium-term (half-year) mortality rate amongst seriously ill hospitalized patients and improving clinical decision making (Connors et al., 1995). With a relatively large sample size of 9,105 and detailed documentation of clinical variables, the SUPPORT dataset allows detection of potential interactions using the strategy implemented in `sail`. We applied `sail` to test for non-linear interactions between acute renal failure or multiple organ system failure (ARF/MOSF), an important predictor for survival rate, and 13 other variables that were deemed clinically relevant. These variables included the number of comorbidities (excluding ARF/MOSF), age, sex, as well as multiple physiological and blood biochemical indices. The response was whether a patient survived after six months since hospitalization.

A total of 8,873 samples had complete data on all variables of interest. We randomly divided these samples into equal sized training/validation/test splits and ran `lassoBT, GLinternet`, and the weak heredity `sail` with cubic B-splines and *α* = 0.1 (as was done in the Nurse Family Partnership program case study). A binomial distribution family was specified for `GLinternet`, whereas `lassoBT` had the same default settings as the simulation study since it did not support a specialized implementation for binary outcomes. We again ran each method on the training data, determined the optimal tuning parameter on the validation data based on the area under the receiver operating characteristic curve (AUC), and assessed AUC on the test data. We repeated this process 200 times and report the results in Table 3.

We found that `sail` achieved similar prediction accuracy to `lassoBT` and `GLinternet`. However, the predictive performance of `lassoBT` and `GLinternet` relied on models which included many more variables. In Figure 4, we visualize the two strongest interaction effects associated with the number of comorbidities and age, respectively. For those having undergone ARF/MOSF, an increased number of comorbidities decreases their chance of survival, while there seems to be no such relationship for non-ARF/MOSF patients. The interaction between ARF/MOSF and age shows the risk incurred by ARF/MOSF is most distinguishing among patients between the ages of 70 and 80.

**Figure 4:**
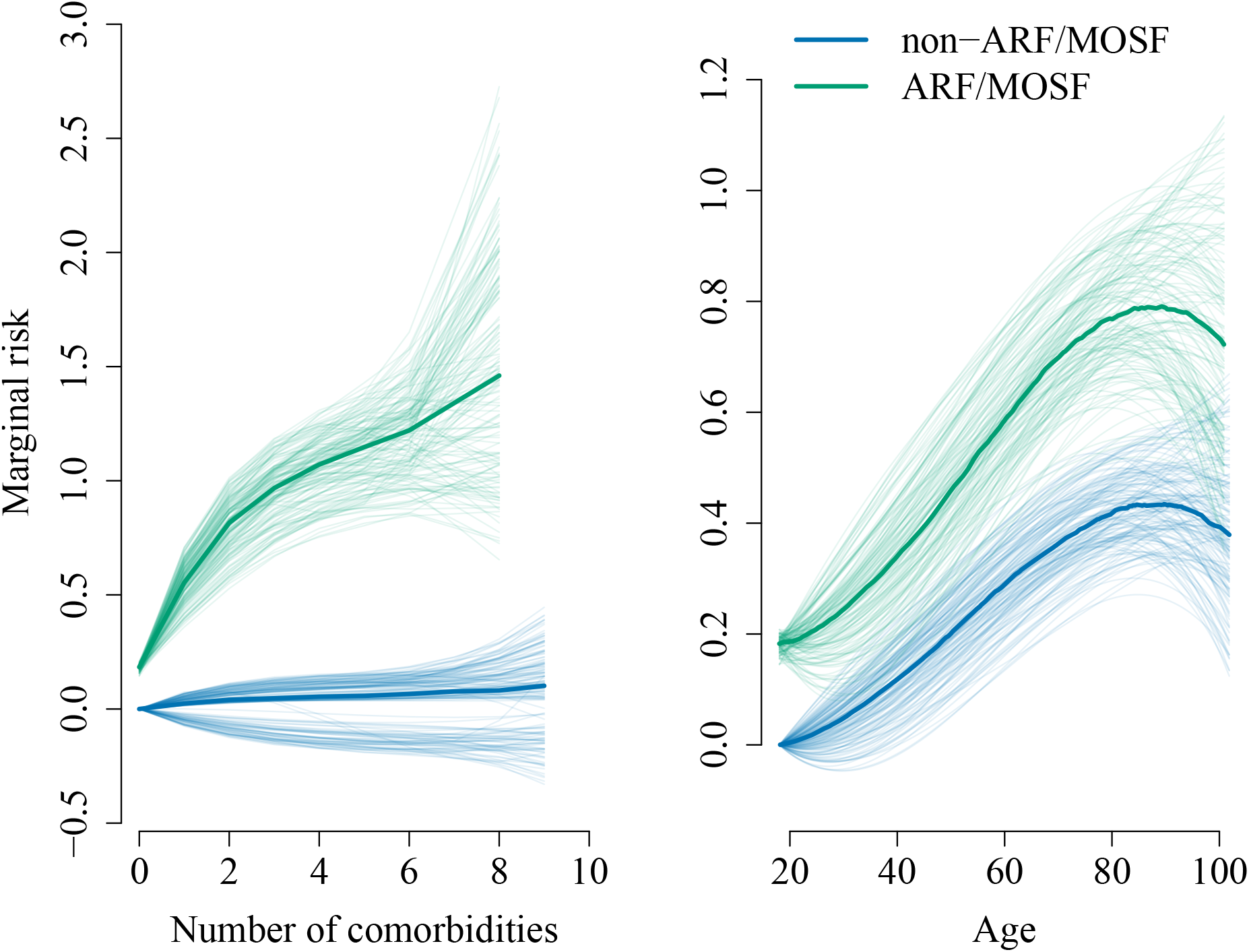
Illustration of estimated interaction effects identified by `sail` for the SUPPORT data. Median prediction curves in dark colors based on 200 train/validate/test splits represent the estimated marginal interaction effects. Coefficients estimated in each of the 200 train/validate/test splits were used to generate prediction curves representing a 90% confidence interval colored in corresponding light colors.

## 6. Discussion

In this article we have introduced the sparse additive interaction learning model `sail` for detecting non-linear interactions with a key environmental or exposure variable in high-dimensional settings. Using a simple reparametrization, we are able to achieve either the weak or strong heredity property without using a complex penalty function. We developed a blockwise coordinate descent algorithm to solve the `sail` objective function for the least-squares loss. We further studied the asymptotic properties of our method and showed that under certain conditions, it possesses the oracle property. All our algorithms have been implemented in a computationally efficient, well-documented and freely available `R` package on CRAN. Furthermore, our method is flexible enough to handle any type of basis expansion including the identity map, which allows for linear interactions. Our implementation allows the user to selectively apply the basis expansions to the predictors, allowing for example, a combination of continuous and categorical predictors. An extensive simulation study shows that `sail, adaptive sail` and `sail weak` outperform existing penalized regression methods in terms of prediction accuracy, sensitivity and specificity when there are non-linear main effects only, as well as interactions with an exposure variable. We then demonstrated the utility of our method to identify non-linear interactions in both biological and epidemiological data. In the NFP program, we showed that individuals who are genetically predisposed to lower educational attainment are those who stand to benefit the most from the intervention. Analysis of the SUPPORT data revealed that those having undergone ARF/MOSF, an increased number of comorbidities decreased their chances of survival, while there seemed to be no such relationship for non-ARF/MOSF patients. In a bootstrap analysis of both datasets, we observed that `sail`tended to select sparser models while maintaining similar prediction performance compared to other interaction selection methods.

Our method however does have its limitations. `sail` can currently only handle *X_E_* · *f*(*X*) or *f*(*X_E_*) · *X* and does not allow for *f*(*X,X_E_*), i.e., only one of the variables in the interaction can have a non-linear effect and we do not consider the tensor product. The reparametrization leads to a non-convex optimization problem which makes convergence rates difficult to assess, though we did not experience any major convergence issues in our simulations and real data analysis. The memory footprint can also be an issue depending on the degree of the basis expansion and the number of variables. Furthermore, the functional form of the covariate effects is treated as known in our method. Being able to automatically select for example, linear vs. nonlinear components, is currently an active area of research in main effects models (Haris et al., 2019). To our knowledge, our proposal is the first to allow for non-linear interactions with a key exposure variable following the weak or strong heredity property in high-dimensional settings. We also provide a first software implementation for these models.

## Acknowledgments

SRB and CMTG were supported by the Ludmer Centre for Neuroinformatics and Mental Health and the Canadian Institutes for Health Research PJT 148620. SRB acknowledges the support of the Natural Sciences and Engineering Research Council of Canada (NSERC), RGPIN-2020-05133. This research was enabled in part by support provided by Calcul Québec (www.calculquebec.ca) and Compute Canada (www.computecanada.ca). The funders had no role in study design, data collection and analysis, decision to publish, or preparation of the manuscript.

## A. Proofs

As shown in the main text, we simplified the notation to make the proofs easier to follow. We summarize the original notation and the corresponding simplified notation in Table 4. This notation then allows us to write down the `sail` estimates as

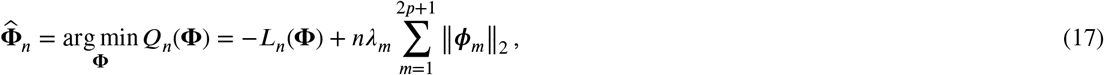

**Table 4.** Correspondence between parameters used to simplify the notation in the proofs. The first row shows the actual parameters used in the loss function. The second row shows the corresponding parameters in the simplified notation. The third row shows the actual tuning parameters used in the penalty function. The fourth row shows the corresponding tuning parameters in the simplified notation. This correspondence greatly simplifies the notation used in the proofs.

|  |  |  |  |  |  |  |  |  |
| --- | --- | --- | --- | --- | --- | --- | --- | --- |
| $\beta_E^*$ | $\theta_1^{*\top}$ | $\theta_2^{*\top}$ | … | $\theta_p^{*\top}$ | $\gamma_{1E}^*$ | $\gamma_{2E}^*$ | … | $\gamma_{pE}^*$ |
| $\phi_1^{*\top}$ | $\phi_2^{*\top}$ | $\phi_3^{*\top}$ | … | $\phi_{p+1}^{*\top}$ | $\phi_{p+2}^{*\top}$ | $\phi_{p+3}^{*\top}$ | … | $\phi_{2p+1}^{*\top}$ |
| $\lambda(1-\alpha)w_E$ | $\lambda(1-\alpha)w_2$ | $\lambda(1-\alpha)w_3$ | … | $\lambda(1-\alpha)w_{p+1}$ | $\lambda\alpha w_{p+2,E}$ | $\lambda\alpha w_{p+3,E}$ | … | $\lambda\alpha w_{2p+1,E}$ |
| $\lambda_1$ | $\lambda_2$ | $\lambda_3$ | … | $\lambda_{p+1}$ | $\lambda_{p+2}$ | $\lambda_{p+3}$ | … | $\lambda_{2p+1}$ |

### A.1. Regularity Conditions

**(C1)** The observation {**V**_*i*_: *i* = 1,…, *n*} are independent and identically distributed with a probability density *f*(**V, Φ**), which has a common support. We assume the density *f* satisfies the following equations:

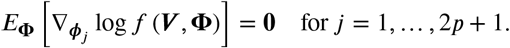

and

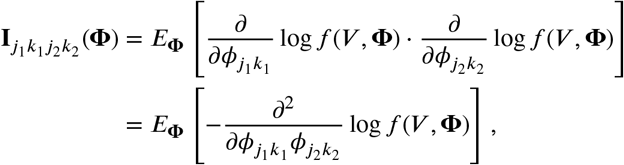

for any *j*_1_, *j*_2_ = 1…, 2*p* + 1, and *k*_1_ = 1,…, *p*_*j*1_, *k*_2_ = 1…, *p*_*j*2_, where *j*_1_, *j*_2_ are the index of group, *k*_1_, *k*_2_ be the index of elements within the corresponding group, *p*_*j*_1__, *p*_*j*_2__ are the group size of *j*_1_, *j*_2_ respectively.
**(C2)** The Fisher information matrix

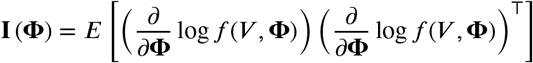

is finite and positive definite at **Φ** = **Φ**\*.
**(C3)** There exists an open set *ω* of Ω that contains the true parameter point **Φ**\* such that for almost all **V** the density *f*(**V, Φ**) admits all third derivatives 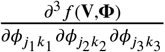 for all **Φ** in *ω* and any *j*_1_, *j*_2_, *j*_3_ = 1,…, 2*p* + 1, and *k*_1_ = 1,…, *p*_*j*1_, *k*_2_ = 1,…, *p*_*j*2_ and *k*_3_ = 1,…, *p*_*j*3_. Furthermore, there exist functions *M*_*j*_1_*k*_1_*j*_2_*k*_2_*j*_3_*k*_3__ such that

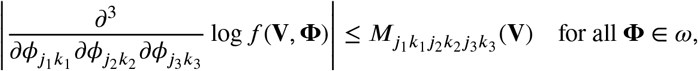

and *m*_*j*_1_*k*_1_*j*_2_*k*_2_*j*_3_*k*_3__ = *E*_**Φ**\*_ [*m*_*j*_1_*k*_1_*j*_2_*k*_2_*j*_3_*k*_3__ (**V**)] < ∞.

### A.2. Lemma 1 proof

Let 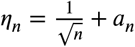 and {**Φ**\* + *η_n_**δ***: ∥***δ***∥_2_ ≤ *C*} be the ball around **Φ**\* for 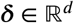, where *d* is the dimension of the design matrix and *C* is some constant. Under the regularity assumptions, we show that there exists a local minimizer 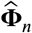 of *Q_n_*(**Φ**) such that 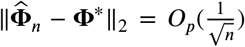. For this proof, we adopt the approaches outlined in (Fan and Li, 2001; Choi et al., 2010; Nardi et al., 2008; Wang et al., 2007) and extend it to our situation. Let 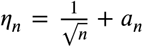 and {**Φ**\* + *η_n_**δ***: ∥***δ***∥_2_ ≤ *C*} be the ball around **Φ**\* for 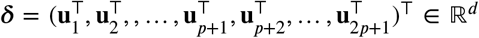, where *d* is the dimension of the design matrix and *C* is some constant. The objective function is given by

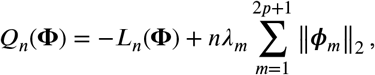

Define

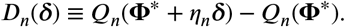

Then for ***δ*** that satisfies ∥***δ***∥_2_ = *C*, we have

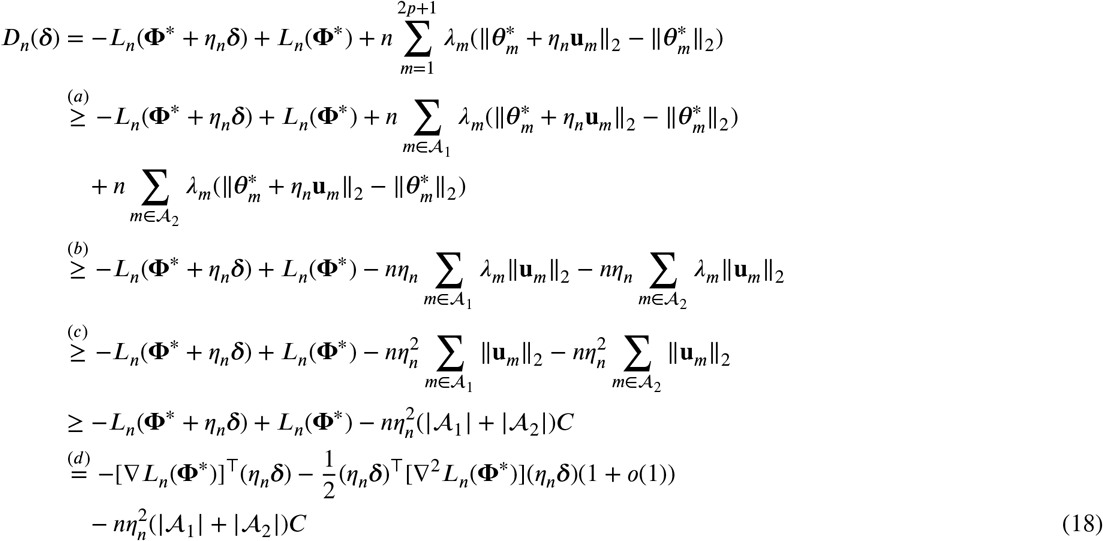

Inequality (a) is by the fact that 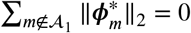 and 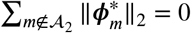. Inequality (b) is due to the reverse triangle inequality ∥*a*∥_2_ – ∥*b*∥_2_ ≥ – ∥*a* – *b*∥_2_. Inequality (c) is by *λ_m_* ≤ *a_n_* ≤ *η_n_* for 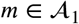 and 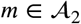. Equality (d) is by the standard argument on the Taylor expansion of the loss function:

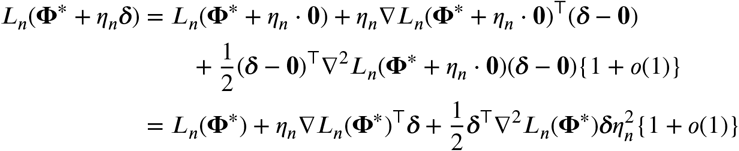

We split (18) into three parts:

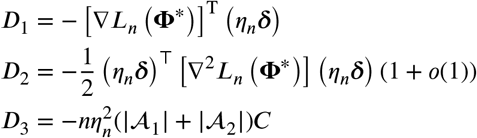

Then

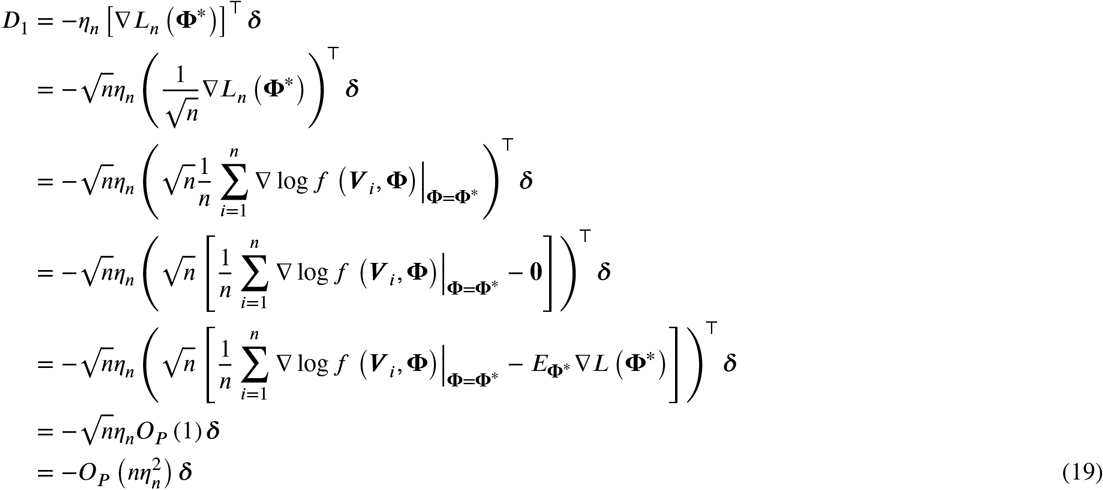

The last equation is by 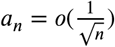 and

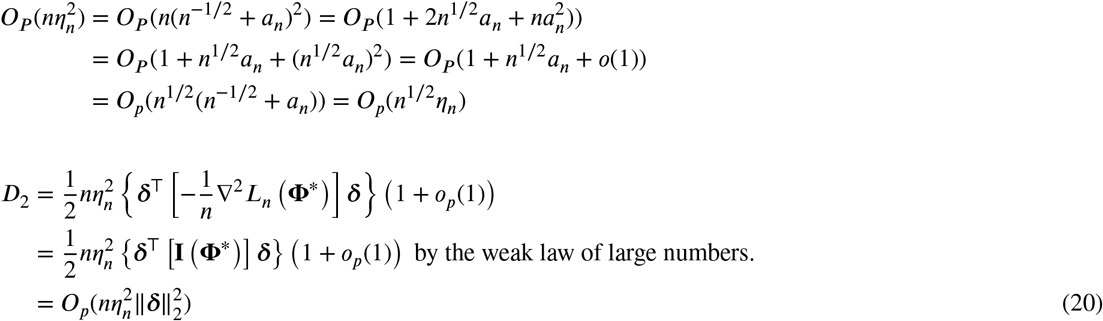

Combining (19) and (20) with (18) gives:

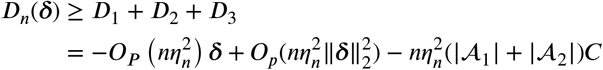

We can see that the first term *D*_1_ is linear in ***δ*** and the second term *D*_2_ is quadratic in ***δ***. We can conclude that for a large enough constant *C* = ∥***δ***∥_2_, *D*_2_ dominates *D*_1_ and *D*_3_. Note that this is a positive term since *I*(**Φ**) is positive definite at **Φ** = **Φ**\* by regularity condition (C2). Therefore, for each *ε* > 0, there exists a large enough constant *C* such that, for large enough *n*

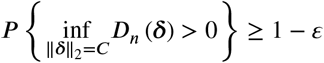

This implies with probability at least 1 − *ε* that the empirical likelihood *Q_n_* has a local minimizer in the ball {**Φ**\* + *η_n_**δ***: ∥***δ***∥_2_ ≤ *C*} (since *Q_n_* is bounded and {**Φ**\* + *α_n_**δ***: ∥***δ***∥_2_ ≤ *C*} is closed). In other words, there exists a local solution 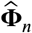 such that 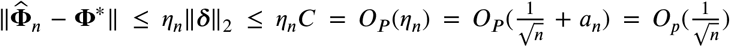, since 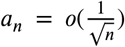. Hence, 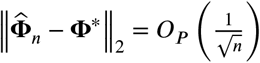.

### A.3. Theorem 1 proof

We first consider consistency for the main effects 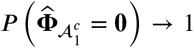. Following (Fan and Li, 2001; Choi et al., 2010), it is sufficient to show that for all 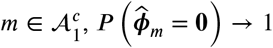, which implies that 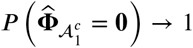, i.e., the 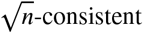 estimate 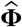 has oracle property 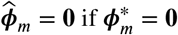. Denote

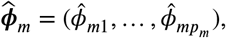

where *p_m_* is the group size of 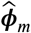. Let 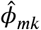 be the *k*-th entry of 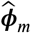. Note that if 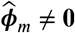, then 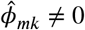 for *k* = 1,…, *p_m_*, then penalty function 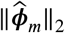 becomes differentiable. Therefore *ϕ_mk_* for *k* = 1,…, *p_m_* must satisfy the following normal equation

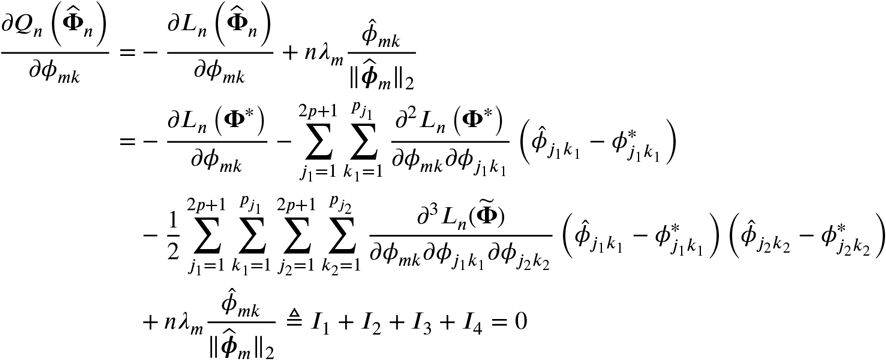

where 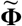 lies between 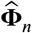 and **Φ**\*. By the regularity conditions and Lemma (1) that 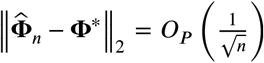, the first term is of the order 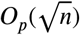

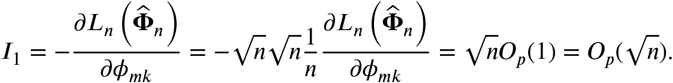

Then the second is of the order 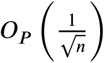 and the third term is of the order 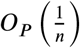. Hence

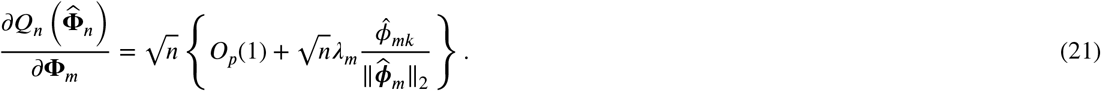

As 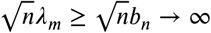 for 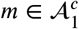 from the assumption, therefore we know that *I*_4_ dominates *I*_1_, *I*_2_ and *I*_3_ in (21) with probability tending to one. This means that (21) cannot be true as long as the sample size is sufficiently large. As a result, we can conclude that with probability tending to one, the estimate 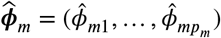 must be in a position where 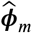 is not differentiable. Hence 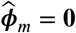 for all 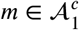. Hence 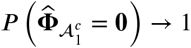. This completes the proof.

Next, we prove that for the interactions 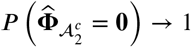. For 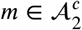 s.t. 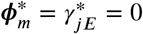 but *β_E_* ≠ 0 and 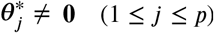, we can prove 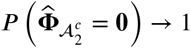 by a similar reasoning, which further implies that 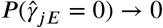. For 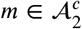 such that 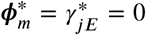 and either *β_E_* = 0 or 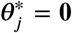 (1 ≤ *j* ≤ *p*): without loss of generality, assume that 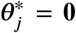. Notice that 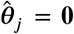 implies 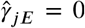, since if 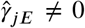, the value of the loss function does not change but the value of the penalty function will increase. Because we already prove 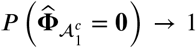, therefore we get 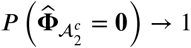 as well for this case.

### A.4. Theorem 2 proof

By Lemma 1 and Theorem 1, there exists a 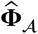 that is a 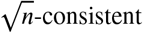 local minimizer of 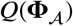, therefore 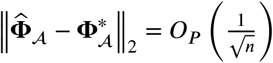 and 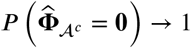. Thus satisfies (with probability tending to 1):

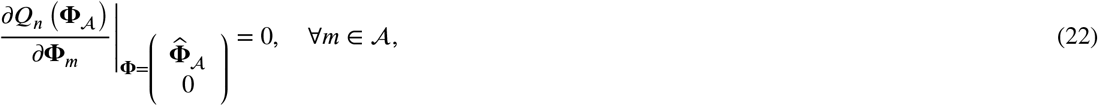

that is

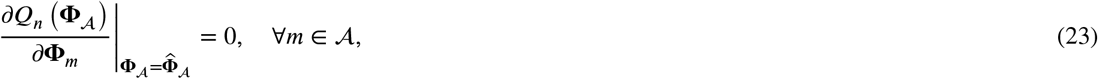

where

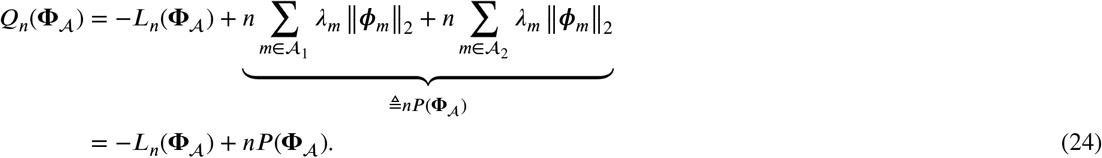

From (23) and (24) we have

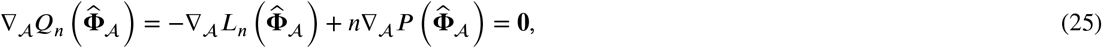

with probability tending to 1.

Denote **Σ** = diag{*o_p_*(1),…, *o_p_*(1)}. We then expand 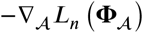 at 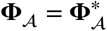 in (25):

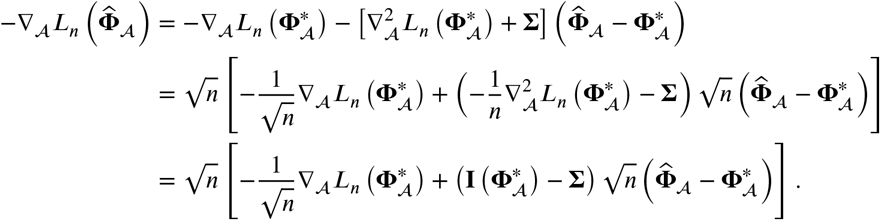

The third line follows by

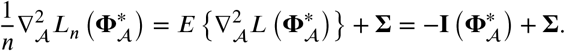

Denote

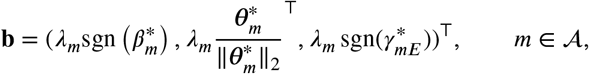

We also expand 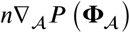 at 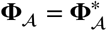 in (25):

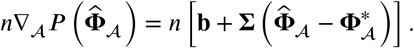

And due to the fact that 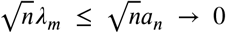 for 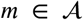 and 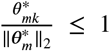 for any 1 ≤ *k* ≤ *p_m_*, we know that 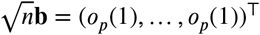 Thus,

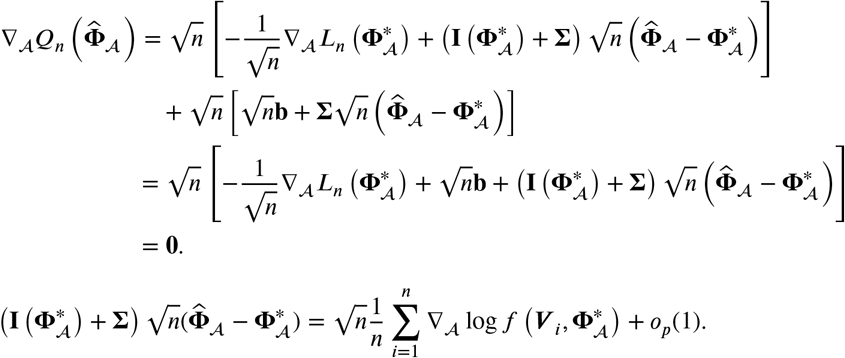

Therefore, by the central limit theorem, we know that

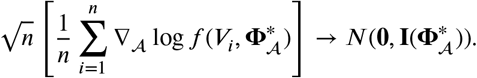

Hence,

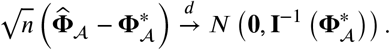

## B. Algorithm Details

In this section we provide more specific details about the algorithms used to solve the `sail` objective function. We assume that *Y*, **Ψ***_j_*, *X_E_* and *X_E_*◦**ψ***_j_* have been centered by their sample means 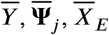, and 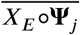, respectively. Here, 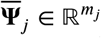 and 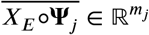 represent the column means of **Ψ***_j_* and *X_E_*◦**ψ***_j_*, respectively. Since the intercept (*β*_0_) is not penalized and all variables have been centered, we can omit it from the loss function and compute it once the algorithm has converged for all other parameters. The strong heredity `sail` model with least-squares loss has the form

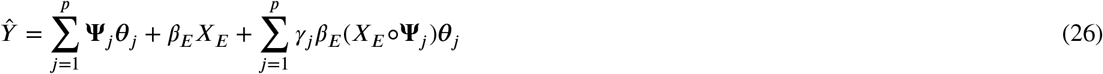

and the objective function is given by

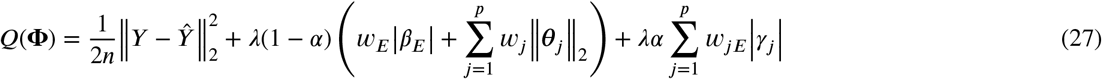

Solving (27) in a blockwise manner allows us to leverage computationally fast algorithms for *ℓ*_1_ and *ℓ*_2_ norm penalized regression. Denote the *n*-dimensional residual column vector 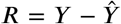. The subgradient equations are given by

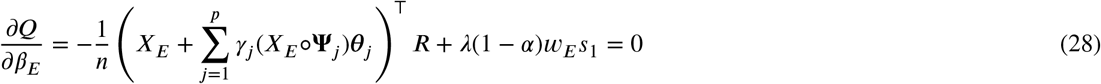

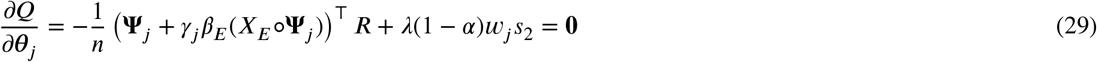

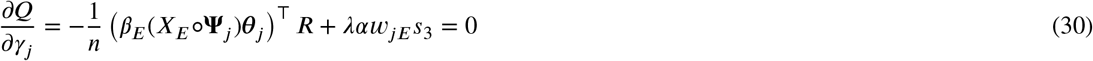

where *s*_1_ is in the subgradient of the *ℓ*_1_ norm:

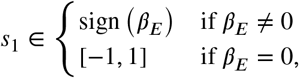

*s*_2_ is in the subgradient of the *ℓ*_2_ norm:

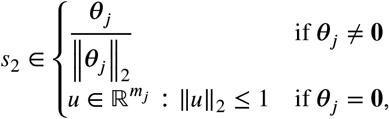

and *s*_3_ is in the subgradient of the *ℓ*_1_ norm:

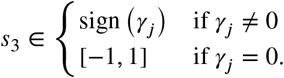

Define the partial residuals, without the *j*th predictor for *j* = 1,…, *p*, as

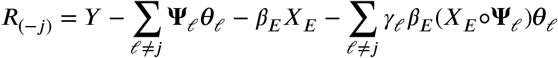

the partial residual without *X_E_* as

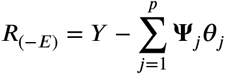

and the partial residual without the *j*th interaction for *j* = 1,…, *p*, as

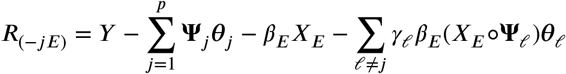

From the subgradient equations (28)–(30) we see that

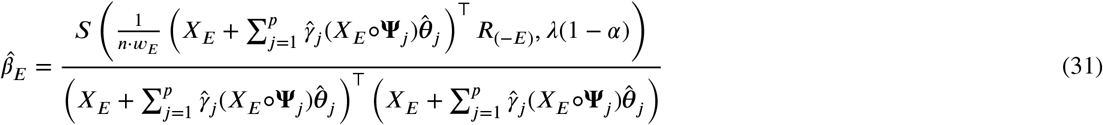

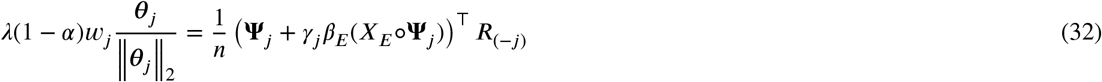

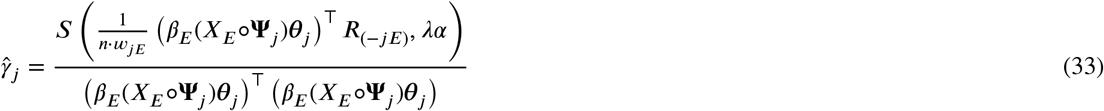

where *S*(*x, t*) = sign(*x*)(|*x*| – *t*) is the soft-thresholding operator. Given these estimates, the intercept can be computed using the following equation:

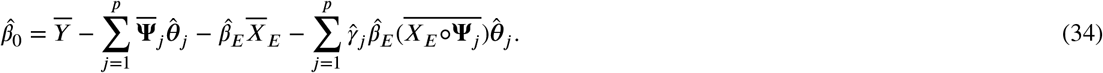

We see from (31) that there is a closed form solution for *β_E_*. From (33), each *γ_j_* also has a closed form solution and can be solved efficiently for *j* = 1,…, *p* using a coordinate descent procedure (Friedman et al., 2010). Since there is no closed form solution for *β_j_*, we use a quadratic majorization technique (Yang and Zou, 2015) to solve (32). Furthermore, we update each ***θ**_j_* in a coordinate wise fashion and leverage this to implement further computational speedups which are detailed in Supplemental Section B.2. From these estimates, we compute the interaction effects using the reparametrizations presented in Table 1, e.g., 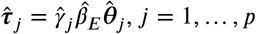 for the strong heredity `sail` model.

### B.1. Least-Squares sail with Strong Heredity

A more detailed algorithm for fitting the least-squares `sail` model with strong heredity is given in Algorithm 3.

#### Algorithm 3 Blockwise Coordinate Descent for Least-Squares sail with Strong Heredity

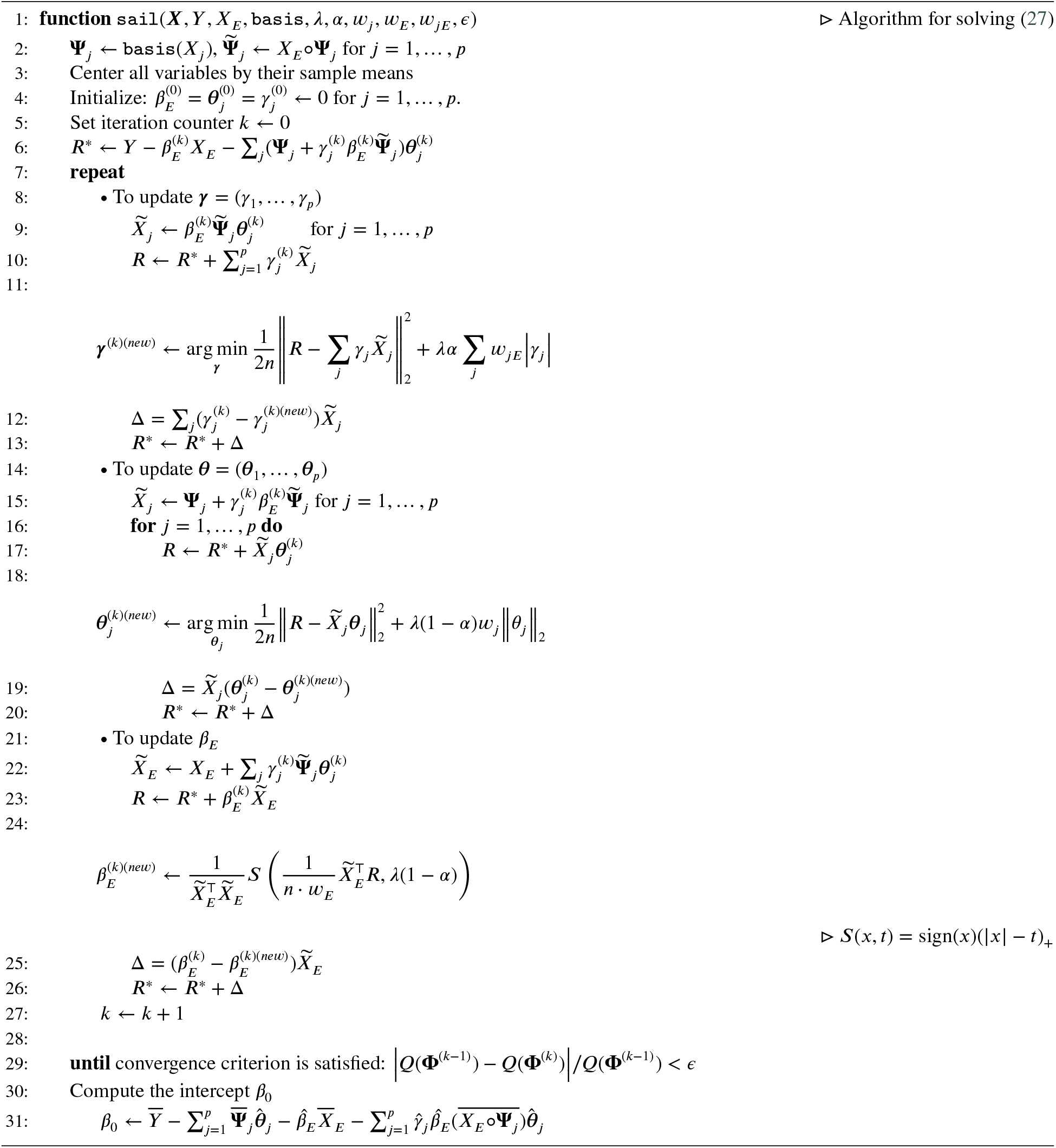

### B.2. Details on Update for *θ*

Here we discuss a computational speedup in the updates for the ***θ*** parameter. The partial residual (*R_s_*) used for updating ***θ**_s_* (*s* ∈ 1,…, *p*) at the *k*th iteration is given by

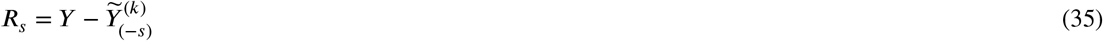

where 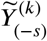 is the fitted value at the *k*th iteration excluding the contribution from **Ψ**_*s*_:

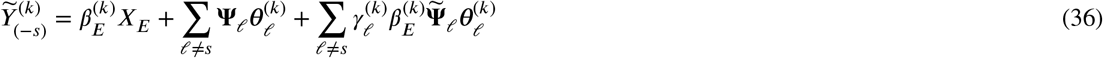

Using (36), (35) can be re-written as

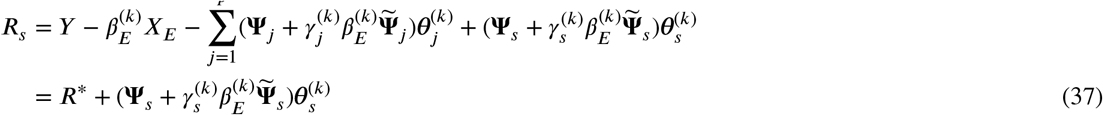

where

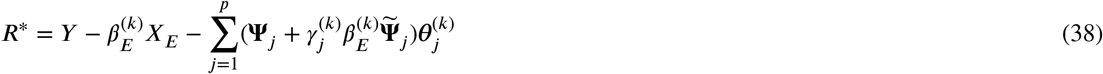

Denote 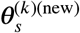 the solution for predictor *s* at the *k*th iteration, given by:

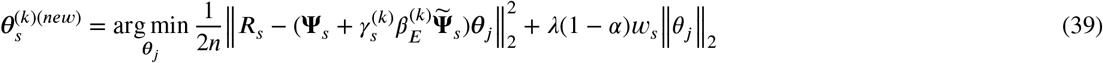

Now we want to update the parameters for the next predictor ***θ***_*s*+1_ (*s* + 1 ∈ 1,…, *p*) at the *k*th iteration. The partial residual used to update ***θ***_*s*+1_ is given by

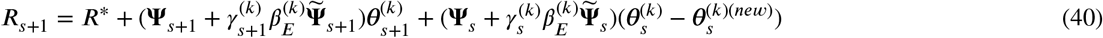

where *R*\* is given by (38), 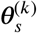 is the parameter value prior to the update, and 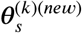 is the updated value given by (39). Taking the difference between (37) and (40) gives

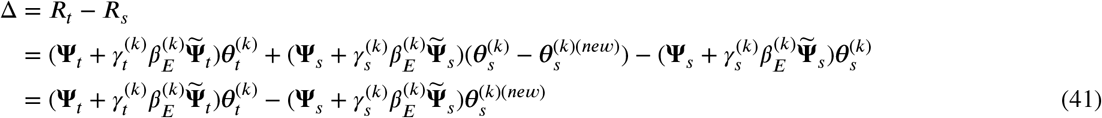

Therefore *R_t_* = *R_s_* + Δ, and the partial residual for updating the next predictor can be computed by updating the previous partial residual by Δ, given by (41). This formulation can lead to computational speedups especially when Δ = 0, meaning the partial residual does not need to be re-calculated.

### B.3. Maximum penalty parameter (*λ_max_*) for strong heredity

The subgradient equations (28)–(30) can be used to determine the largest value of *λ* such that all coefficients are 0. From the subgradient Equation (28), we see that *β_E_* = 0 is a solution if

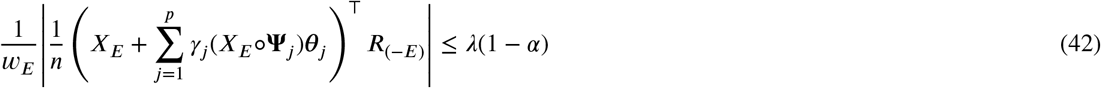

From the subgradient Equation (29), we see that ***θ**_j_* = **0** is a solution if

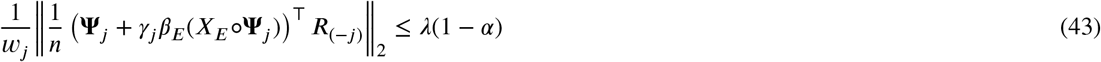

From the subgradient Equation (30), we see that *γ_j_* =0 is a solution if

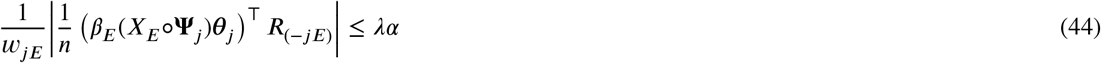

Due to the strong heredity property, the parameter vector (*β_E_*, ***θ***_1_,…, ***θ**_p_*, *γ*_1_,…, *γ_p_*) will be entirely equal to ***θ*** if (*β_E_*, ***θ***_1_,…, ***θ**_p_*) = **0**. Therefore, the smallest value of *λ* for which the entire parameter vector (excluding the intercept) is **0** is:

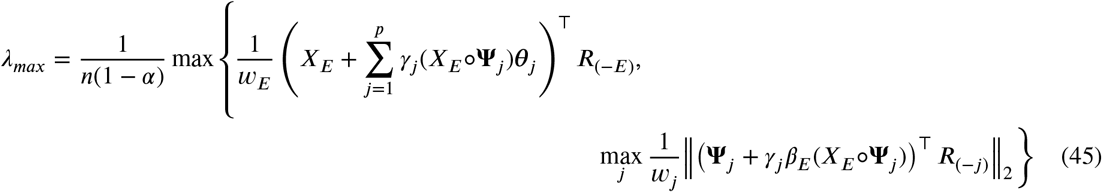

which reduces to

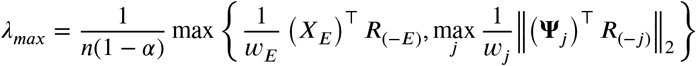

### B.4. Least-Squares sail with Weak Heredity

We assume the same centering constraints as in Section B.1. The least-squares `sail` model with weak heredity has the form

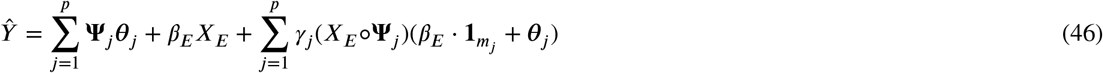

The objective function is given by

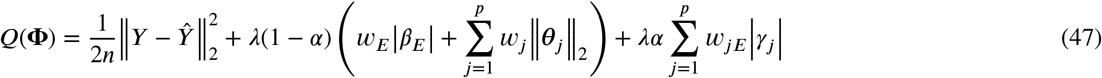

Denote the *n*-dimensional residual column vector 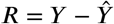. The subgradient equations are given by

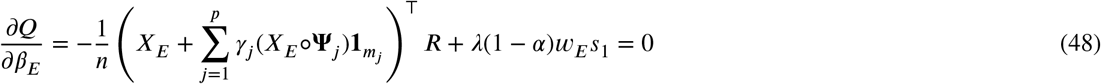

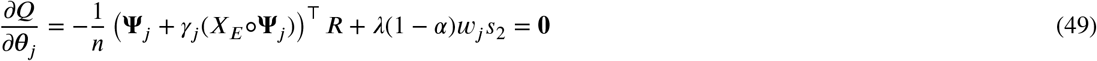

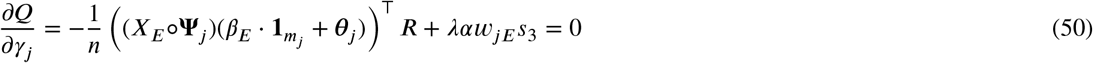

where *s*_1_ is in the subgradient of the *ℓ*_1_ norm:

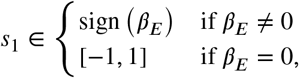

*s*_2_ is in the subgradient of the *ℓ*_2_ norm:

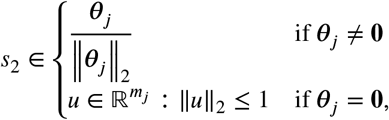

and *s*_3_ is in the subgradient of the *ℓ*_1_ norm:

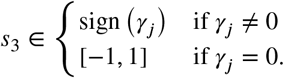

Define the partial residuals, without the *j*th predictor for *j* = 1,…, *p*, as

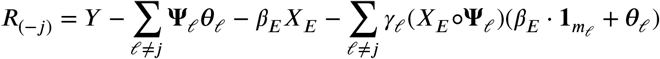

the partial residual without *X_E_* as

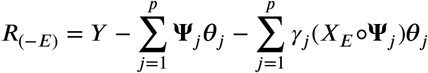

and the partial residual without the *j*th interaction for *j* = 1,…, *p*

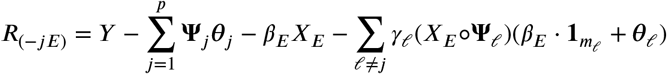

From the subgradient Equation (48), we see that *β_E_* = 0 is a solution if

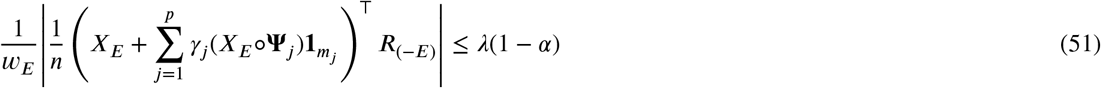

From the subgradient Equation (49), we see that ***θ**_j_* = **0** is a solution if

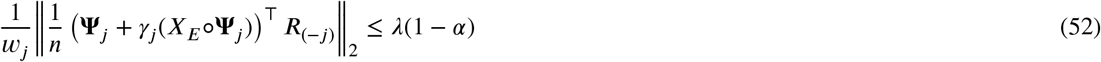

From the subgradient Equation (50), we see that *γ_j_* = 0 is a solution if

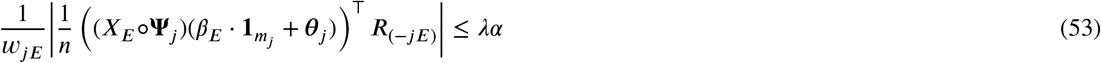

From the subgradient equations we see that

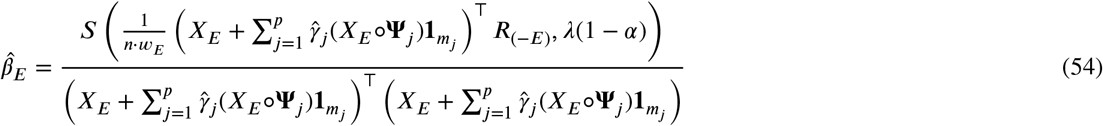

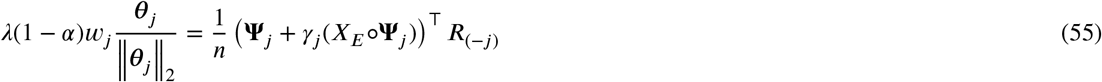

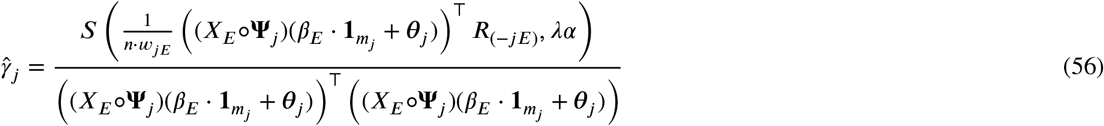

where *S*(*x, t*) = sign(*x*)(|*x*| − *t*) is the soft-thresholding operator. As was the case in the strong heredity `sail` model, there is a closed form solution for *β_E_*, each *γ_j_* also has a closed form solution and can be solved efficiently for *j* = 1,…, *p* using the coordinate descent procedure implemented in the `glmnet` package (Friedman et al., 2010), while we use the quadratic majorization technique implemented in the `gglasso` package (Yang and Zou, 2015) to solve (55). Algorithm 4 details the procedure used to fit the least-squares weak heredity `sail` model.

#### Algorithm 4 Coordinate descent for least-squares sail with weak heredity

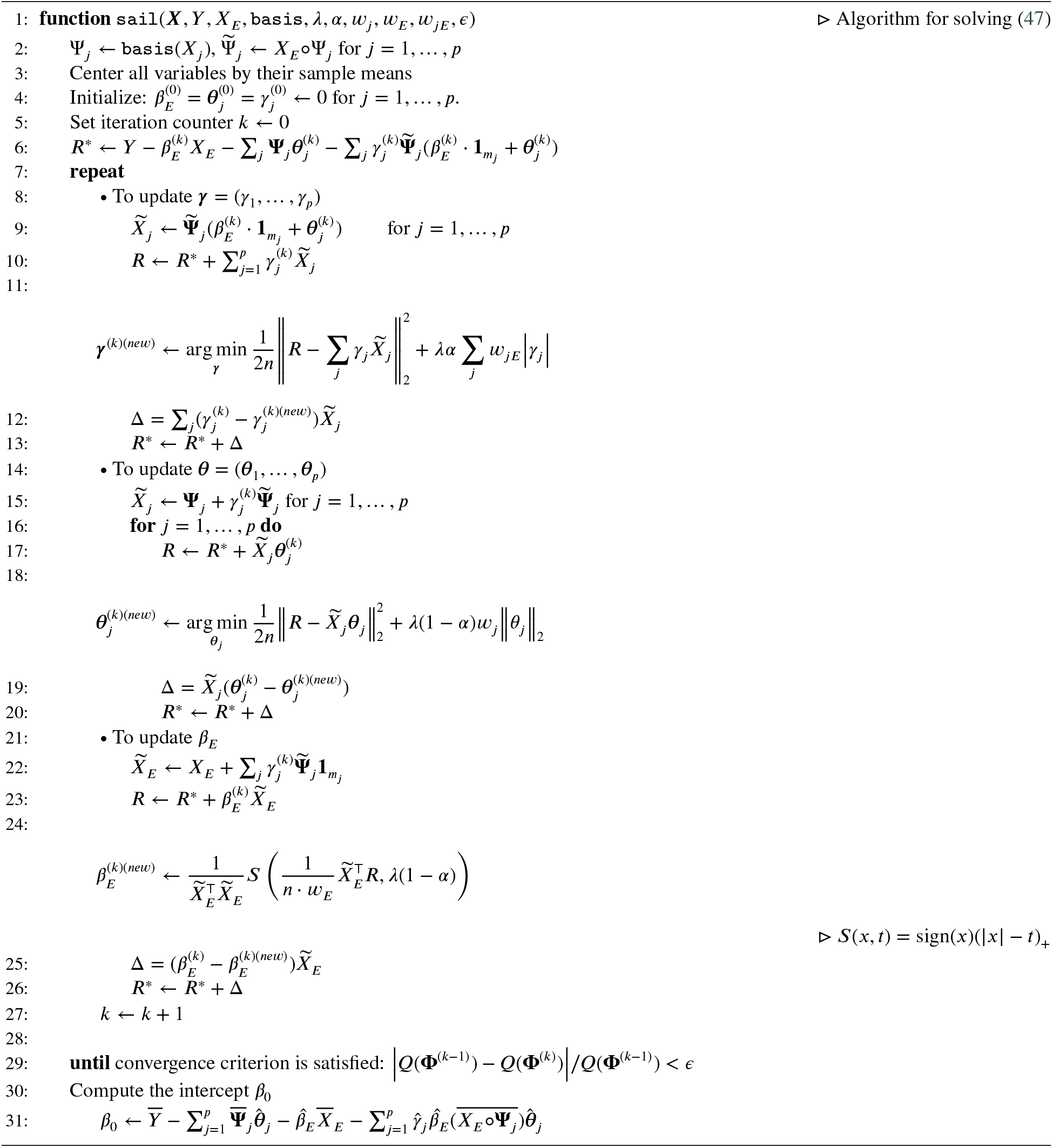

#### B.4.1. Maximum penalty parameter (λ_max_)for weak heredity

The smallest value of *λ* for which the entire parameter vector (*β_E_, **θ***_1_,…, ***θ**_p_*, *γ*_1_,…, *γ_p_*) is **0** is:

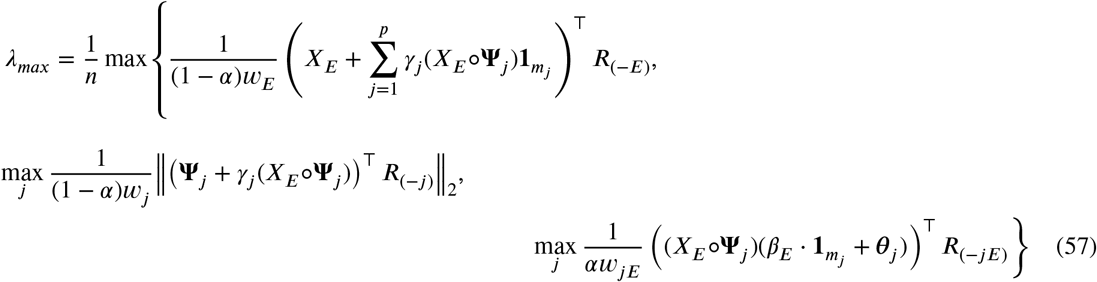

which reduces to

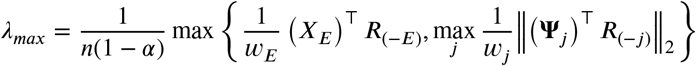

This is the same *λ_max_* as the least-squares strong heredity `sail` model.

## C. Additional Simulation Results

We visually inspected whether our method could correctly capture the shape of the association between the predictors and the response for both main and interaction effects. To do so, we plotted the true and predicted curves for scenario 1a) only. Figure 5 shows each of the four main effects with the estimated curves from each of the 200 simulations along with the true curve. We can see the effect of the penalty on the parameters, i.e., decreasing prediction variance at the cost of increased bias. This is particularly well illustrated in the bottom right panel where `sail` smooths out the very wiggly component function *f*_4_(*x*). Nevertheless, the primary shapes are clearly being captured.

**Figure 5:**
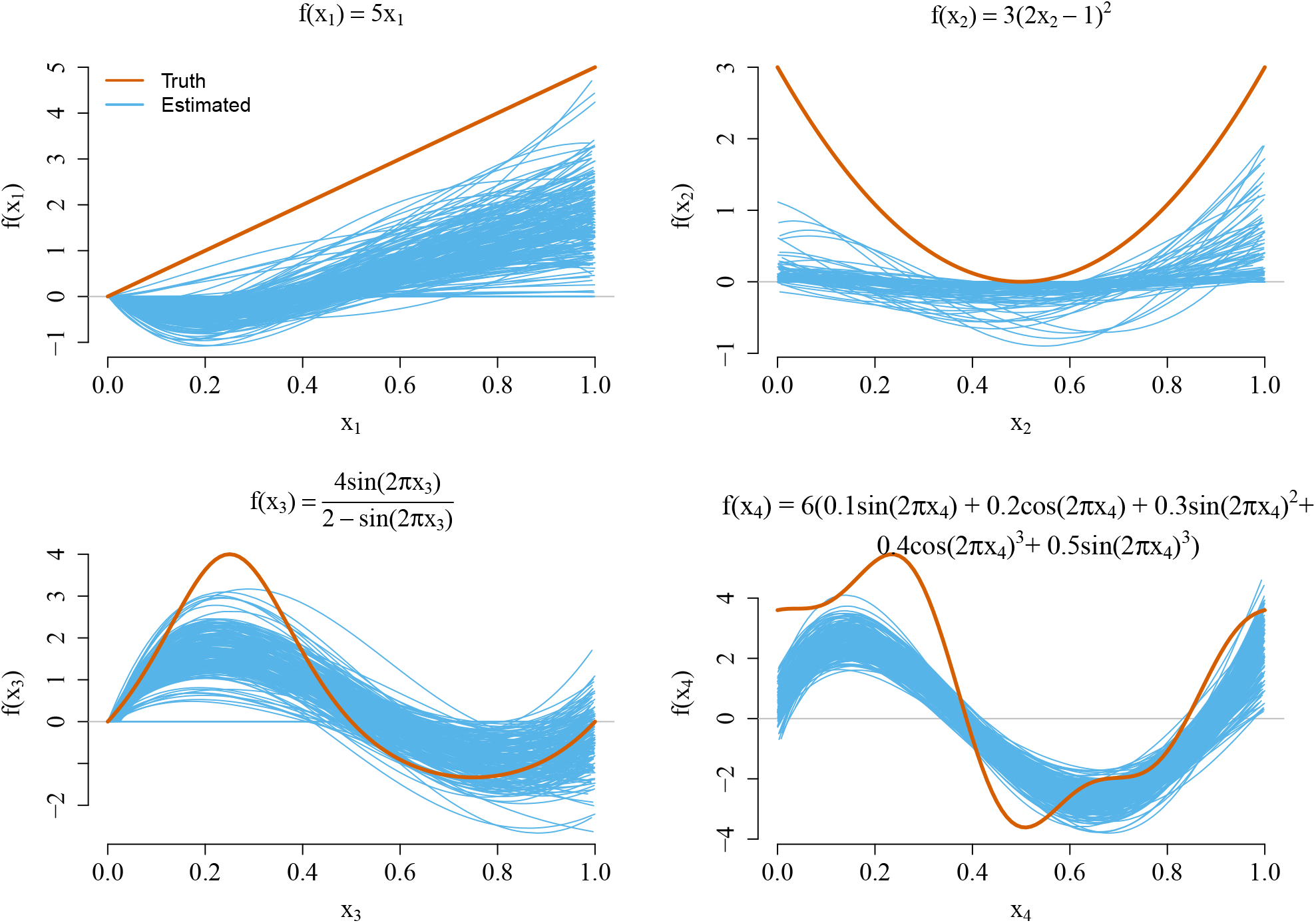
True and estimated main effect component functions for scenario 1a). The estimated curves represent the results from each one of the 200 replications conducted.

To visualize the estimated interaction effects, we ordered the 200 simulation runs by the Euclidean distance between the estimated and true regression functions. Following Radchenko et al. (Radchenko and James, 2010), we then identified the 25th, 50th, and 75th best simulations and plotted, in Figures 6 and 7, the interaction effects of *X_E_* with *f*_3_(*X*_3_) and *f*_4_(*X*_4_), respectively. We see that `sail` does a good job at capturing the true interaction surface for *X_E_* · *f*_3_(*X*_3_). Again, the smoothing and shrinkage effect is apparent when looking at the interaction surfaces for *X_E_* · *f*_4_(*X*_4_).

**Figure 6:**
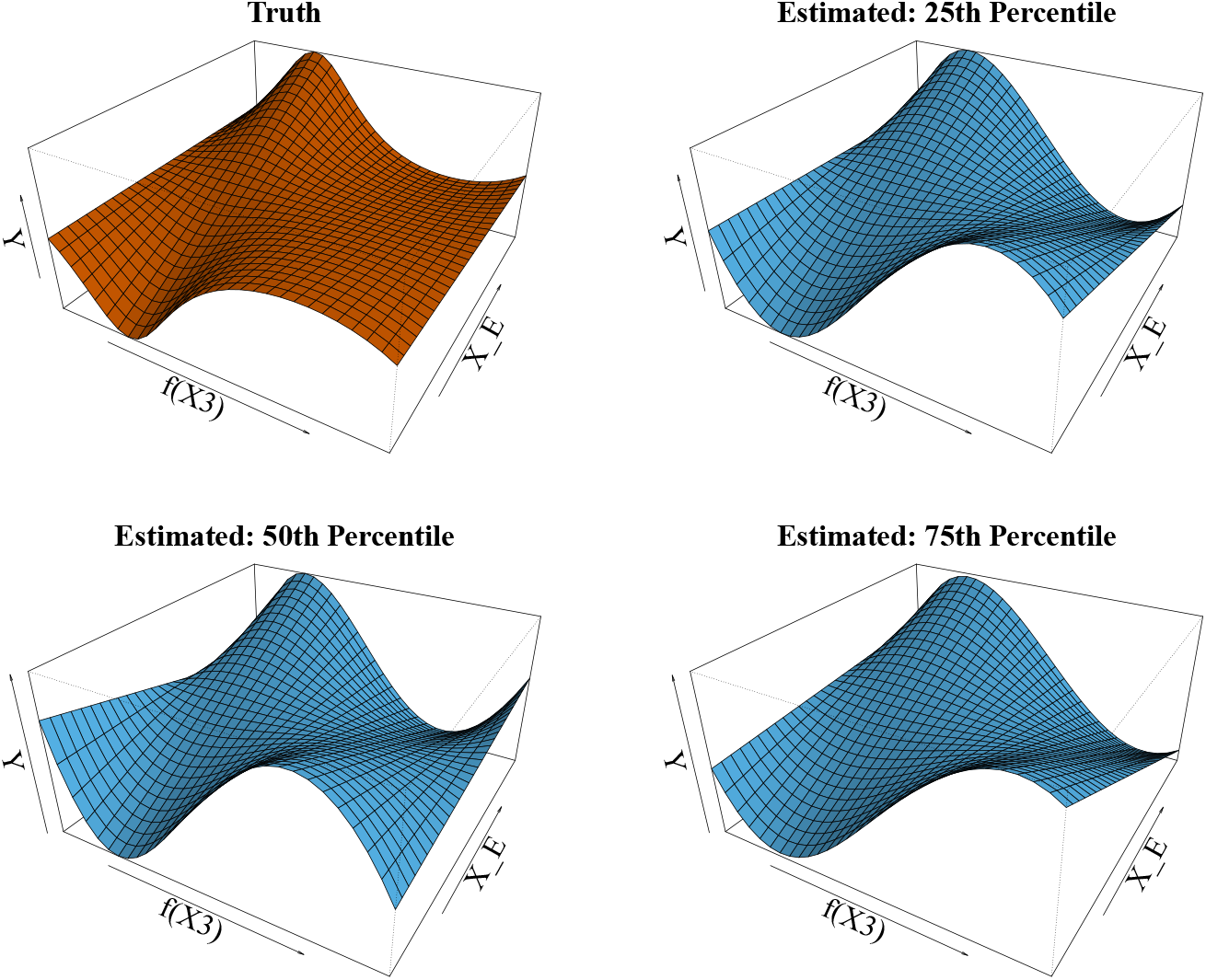
True and estimated interaction effects for *X_E_* · *f*_3_(*X*_3_) in simulation scenario 1a).

**Figure 7:**
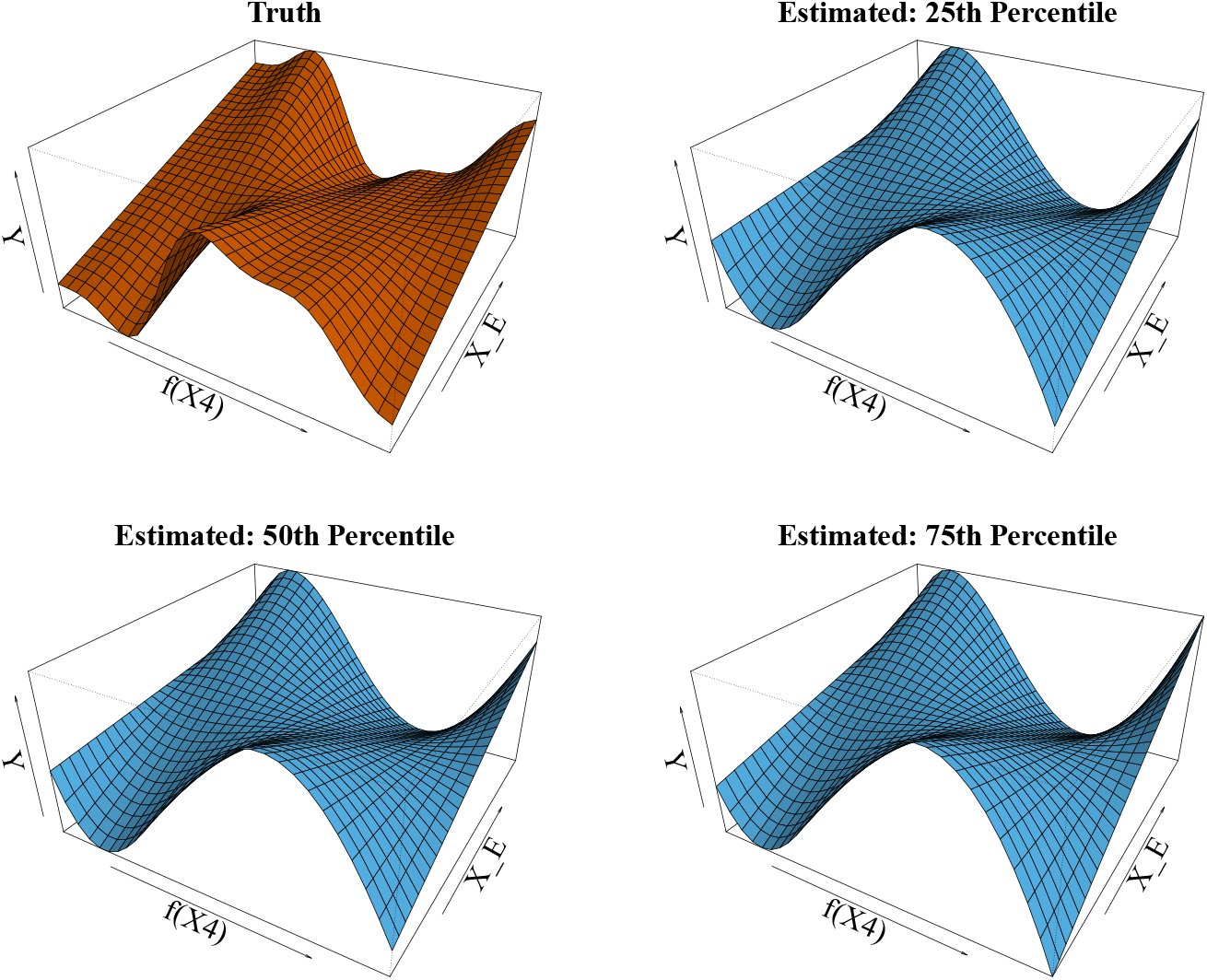
True and estimated interaction effects for *X_E_* · *f*_4_(*X*_4_) in simulation scenario 1a).

In Figure 8 we visualize the variable selection results from 210 replications of the simulation study for strong hierarchy `sail` using UpSet plots (Conway, Lex and Gehlenborg, 2017). Shown are the selected models and their frequencies. We can see that the environment variable is always selected across all simulation scenarios and replications.

**Figure 8:**
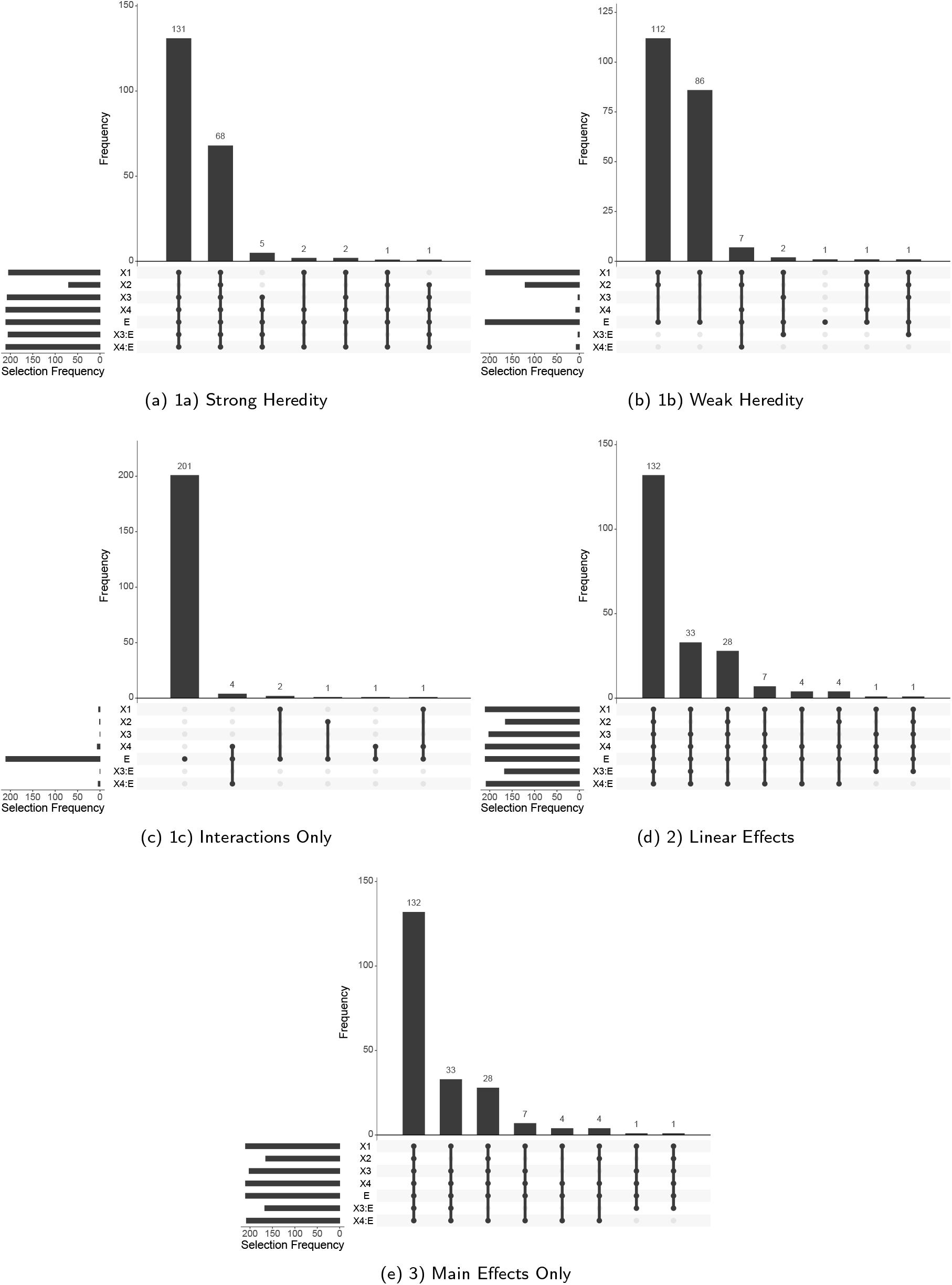
Variable selection results from 210 replications of the simulation study for strong hierarchy `sail` visualized using UpSet plots (Conway et al., 2017). Shown are the selected models and their frequencies. We can see that the environment variable is always selected across all simulation scenarios and replications.

## D. Additional Results on PRS for Educational Attainment

**Figure 9:**
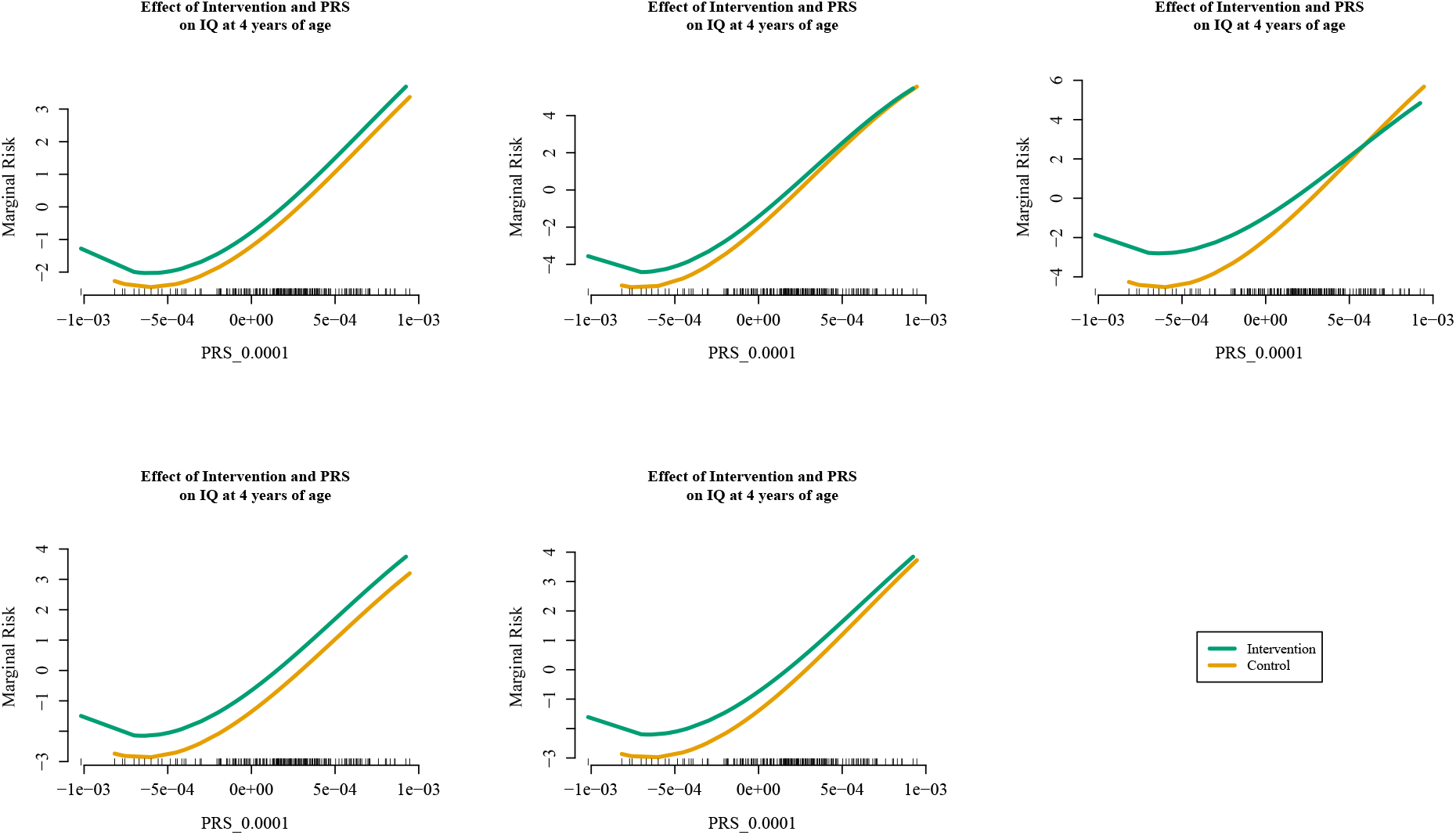
Estimated interaction effect identified by the weak heredity `sail` using cubic B-splines and *α* = 0.1 for the Nurse Family Partnership data for the 5 imputed datasets. Of the 189 subjects, 19 IQ scores were imputed using `mice` (Buuren and Groothuis-Oudshoorn, 2010). The selected model, chosen via 10-fold cross-validation, contained three variables: the main effects for the intervention and the PRS for educational attainment using genetic variants significant at the 0.0001 level, as well as their interaction.

**Figure 10:**
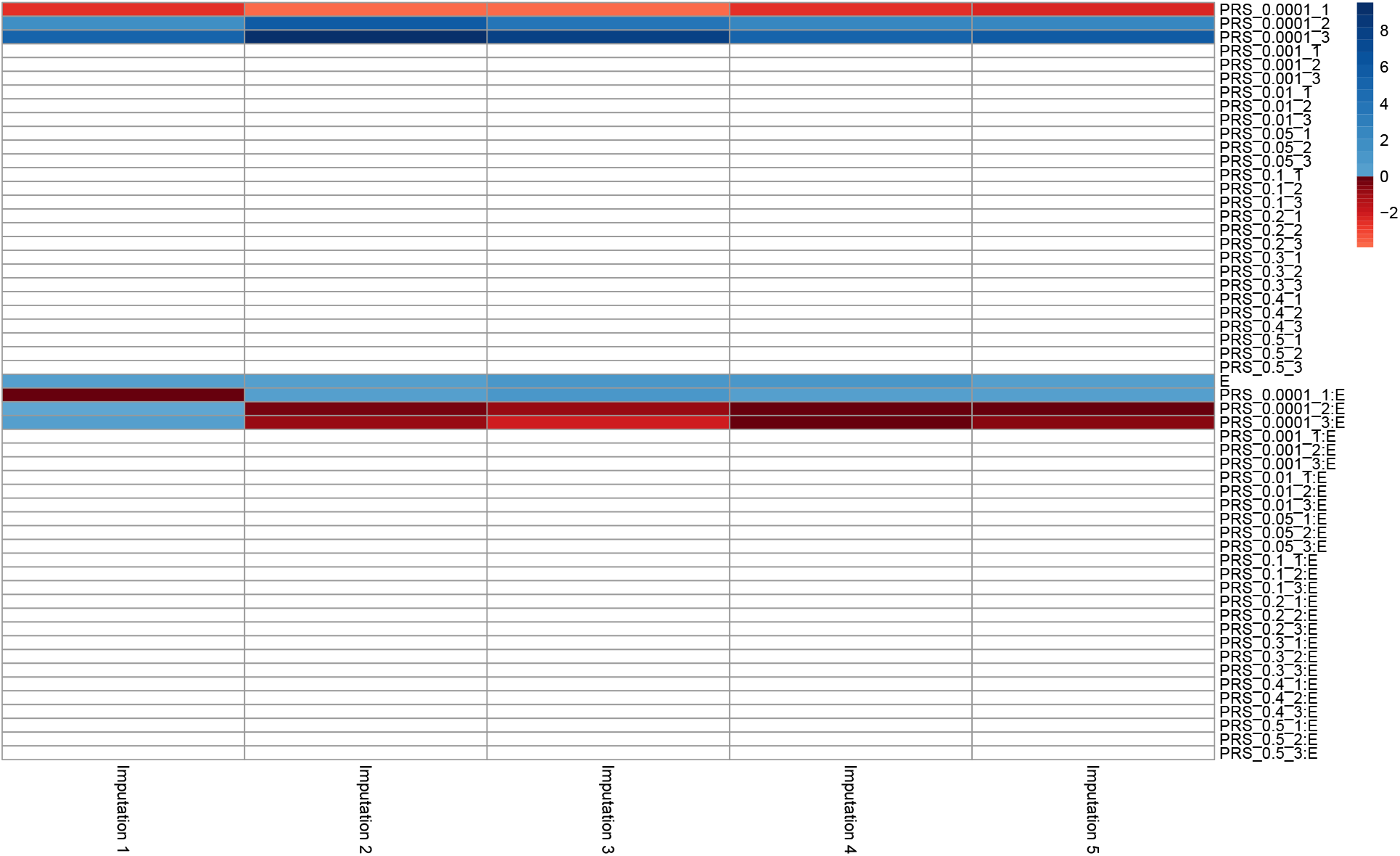
Coefficient estimates obtained by the weak heredity `sail` using cubic B-splines and *α* = 0.1 for the Nurse Family Partnership data for the 5 imputed datasets. Of the 189 subjects, 19 IQ scores were imputed using `mice` (Buuren and Groothuis-Oudshoorn, 2010). The selected model, chosen via 10-fold cross-validation, contained three variables: the main effects for the intervention and the PRS for educational attainment using genetic variants significant at the 0.0001 level, as well as their interaction. This results was consistent across all 5 imputed datasets. The white boxes indicate a coefficient estimate of 0.

## E. Data Availability and Code to Reproduce Results

The R scripts used to simulate the data for the simulation studies in Section 4 are provided along with the code for each of the methods being compared. The data used for the two real data analyses in Section 5 are publicly available. The first dataset from the Nurse Family Partnership program is provided by one of the authors of the manuscript (David Olds). The second dataset from the Study to Understand Prognoses Preferences Outcomes and Risks of Treatment (SUPPORT) is publicly available from the Vanderbilt University Department of Biostatistics website.

### E.1. Datasets

The datasets are available at https://github.com/sahirbhatnagar/sail/tree/master/manuscript/raw_data

1. Nurse Family Partnership program data consists of three files. They are merged together using the script https://github.com/sahirbhatnagar/sail/blob/master/manuscript/bin/PRS_bootstrap.R

- Gen_3PC_scores.txt
- IQ_and_mental_development_variables_for_Sahir_with_study_ID.txt
- NFP_170614_INFO08_nodup_hard09_noambi_GWAS_EduYears_Pooled_beta_withaf_5000pruned_noambi_16Jan2018.score
2. The SUPPORT data consists of a single file:

- https://github.com/sahirbhatnagar/sail/blob/master/manuscript/raw_data/support2.csv

All datasets are in `.txt` format. Code used to read in the datasets are provided in the section below. All output from this project published online is available according to the conditions of the Creative Commons License (https://creativecommons.org/licenses/by-nc-sa/2.0/)

### E.2. Code

The software which implements our algorithm is available in an R package published on CRAN (https://cran.r-project.org/package=sail) version 0.1.0 with MIT license. The paper itself is written in knitr format, and therefore includes both the code and text in the same `.Rnw` file.

The scripts and data used to produce the results in the manuscript are available at https://github.com/sahirbhatnagar/sail/tree/master/manuscript.

The knitr file which contains both the main text and code is available at: https://github.com/sahirbhatnagar/sail/blob/master/manuscript/source/sail_manuscript_v2.Rnw

The manuscript was compiled using R version 3.6.1 with knitr version 1.25.

The bootstrap analysis was run in parallel on a compute cluster with 40 cores. Though this is not necessary to reproduce the results, it definitely speeds up the computation time.

#### E.2.1. Instructions for Use

All tables and figures from the paper can be reproduced by compiling the knitr file. The easiest way to reproduce the results is to download the GitHub repository and compile the knitr file from within an R session as follows:

1. Download the GitHub repository https://github.com/sahirbhatnagar/sail/archive/master.zip
2. From within an R session, run the command: `knitr::knit2pdf`(`‘sail_manuscript_v2.Rnw’`)

Note that to speed up compilation time, we have saved the simulation and bootstrap results in `.RData` files available at https://github.com/sahirbhatnagar/sail/tree/master/manuscript/results. These `.RData` files are called directly by the knitr file.

Note also that the R scripts used to generate the results are called from the knitr file using the ‘code externalization’ functionality of knitr (https://yihui.org/knitr/demo/externalization/). That is, the actual R code is stored in R scripts and not within the knitr file. These R scripts are available at https://github.com/sahirbhatnagar/sail/tree/master/manuscript/bin.

The expected run time to compile the manuscript is about 5 minutes on a standard desktop machine, assuming that you are using the pre-run simulation and bootstrap results.

#### E.2.2. R Package Vignette

A website with two vignettes has been created for our sail package available at https://sahirbhatnagar.com/sail/

The 2 vignettes are:

1. https://sahirbhatnagar.com/sail/articles/introduction-to-sail.html
2. https://sahirbhatnagar.com/sail/articles/user-defined-design.html

